# Molecular signatures associated with successful implantation of the human blastocyst

**DOI:** 10.1101/2023.05.09.539763

**Authors:** Jennifer N. Chousal, Srimeenakshi Srinivasan, Katherine Lee, Cuong To, Kyucheol Cho, Wei Zhang, Ana Lisa Yeo, V. Gabriel Garzo, Mana M. Parast, Louise C. Laurent, Heidi Cook-Andersen

## Abstract

Embryo implantation in humans is remarkably inefficient for reasons that remain largely unexplained, and high rates of implantation failure remain one of the greatest obstacles in treating infertility. The volume of gene expression data available from human embryos has rapidly accumulated in recent years. However, prioritization of these data to identify the subset of genes that determine successful implantation remains a challenge, in part, because comprehensive analyses cannot be performed on the same embryos that are transferred. Here, we leverage clinical morphologic grading—known for decades to correlate with implantation potential—and transcriptome analyses of matched embryonic and abembryonic samples to identify genes and cell-cell interactions enriched and depleted in human blastocysts of good and poor morphology, genome-wide. Unexpectedly, we discovered that the greatest molecular difference was in the state of the extraembryonic primitive endoderm (PrE), with relative deficiencies in PrE development in embryos of poor morphology at the time of embryo transfer. Together, our results support a model in which implantation success is most strongly reflected by factors and signals from the embryonic compartment and suggest that deficiencies in PrE development, in particular, are common among embryos with reduced implantation potential. Our study provides a valuable resource for those investigating the markers and mechanisms of human embryo implantation.

## Introduction

The efficiency of human embryo implantation is remarkably low^1^. Cyclic fecundity rates in fertile women are 20-30%^2, 3^ compared to 30-70% in non-human primates^4^ and 85% in mice^5^. While the precise causes are unknown, high rates of molecular or genetic defects in human embryos are major contributors^6^, leading to a failure to initiate or sustain implantation until pregnancy can be detected clinically. In fact, early implantation failure has been estimated to account for ∼70% of lost pregnancies^2^ and is widely considered to be one of the greatest obstacles in treating infertility. Even in *in vitro* fertilization (IVF) cycles, where embryo quality and ploidy can be assessed before transfer, up to 70% of failed pregnancies are attributed to failed implantation^7^. However, advances in the field have been slow, as the mechanisms critical for successful implantation in humans remain poorly understood and challenging to study. Efforts to identify underlying mechanisms are hindered by multiple technical and ethical limitations in working with human embryos, including the small number of high quality, chromosomally normal embryos available for research, the inability to comprehensively analyze gene expression in the same embryos that are transferred, and the inability to use traditional genetic approaches to knock out factors in specific lineages to establish causal relationships. For these reasons, despite the recent advances in single cell sequencing that have facilitated a rapid expansion in our knowledge of gene expression patterns in the human embryo^8–,17^, a gap remains in connecting these gene expression patterns to functional outcomes in embryo implantation.

Following fertilization, the human embryo undergoes growth and differentiation over ∼7 days to form a blastocyst competent for implantation. Three cell lineages are established in the blastocyst: the trophectoderm (TE), which initiates contact with the endometrium and will form the placenta; the epiblast (EPI), which undergoes gastrulation to form the embryo proper; and the primitive endoderm (PrE), which overlies the EPI and will give rise to the visceral and parietal yolk sacs^8^. The first lineage segregation occurs at Day (D)4-5, when the morula undergoes cavitation to form the blastocyst, separating the outer TE from the inner cell mass (ICM). The timing for EPI and PrE segregation within the ICM remains less established. Lineage marker analyses at the protein level indicate differentiation of EPI and PrE by D6-7^9–12^. RNA profiling suggests that specification begins at earlier stages, with lineage-specific gene signatures for EPI and PrE detectable before D5 and continuing through D5-7^13–16^. Successful lineage specification and differentiation depends not only on innate gene expression programs within each cell type, but also heavily upon cellular cross-talk between each of the cell types^6, 8^. Yet, the full repertoire of these cell-cell interactions, and the consequences of defects in these interactions, has not been delineated. Implantation of the late blastocyst to initiate pregnancy requires that development and differentiation of each lineage has transpired correctly (Fig 1A) and that progression through these events occurs in a timely manner so that the maturation state of the blastocyst lineages and the receptivity of the maternal endometrium complement one another^17, 18^. Accordingly, molecular errors in development that lead to defective or delayed differentiation can result in asynchrony between the embryo stage and endometrium and to failed implantation^17, 18^.

**Fig. 1.**
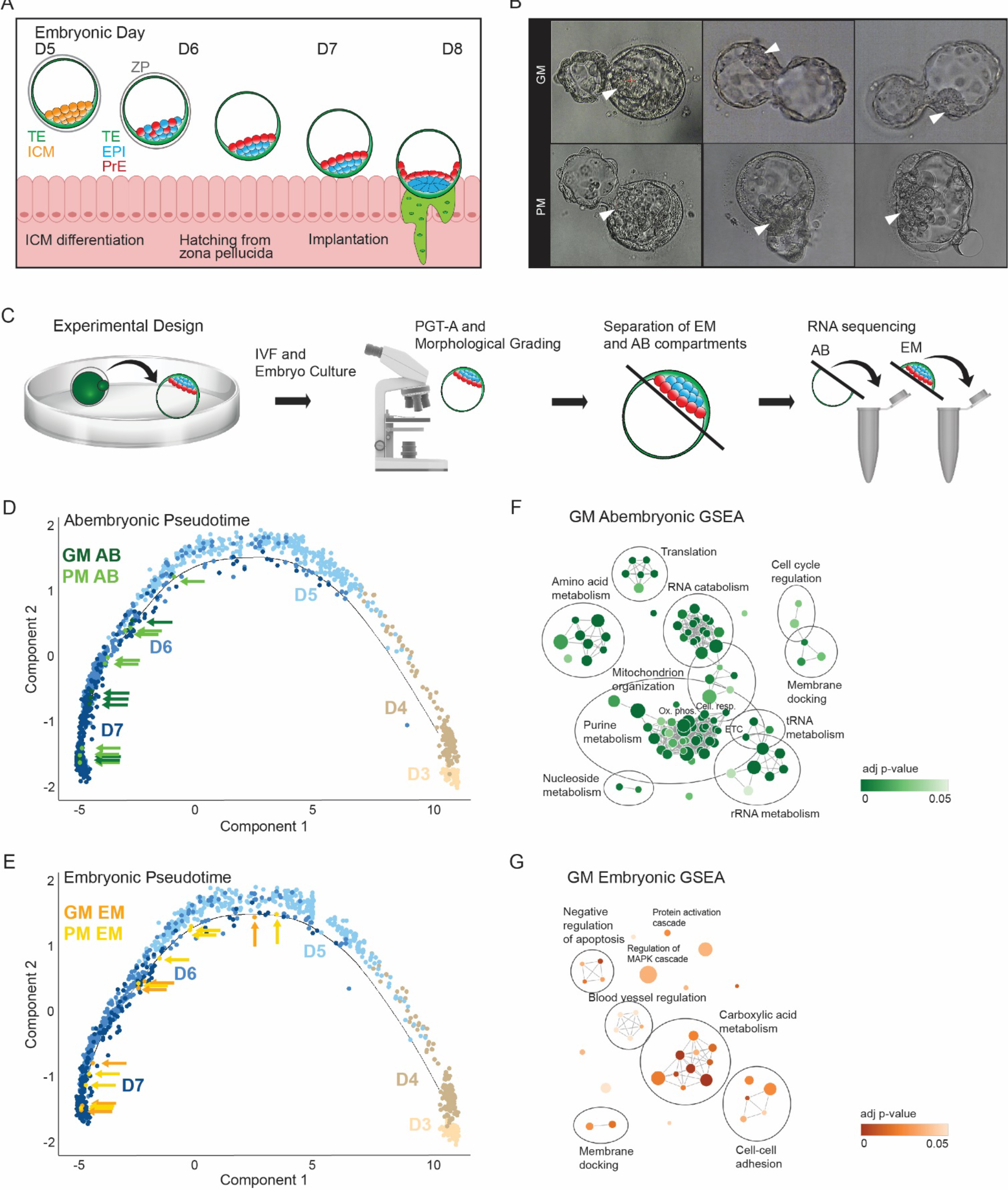
Transcriptome analysis of good and poor morphology human blastocysts. (A) Diagram of human blastocyst maturation and implantation, encompassing embryonic days (D)5-8. At approximately D5-6, the ICM begins to differentiate into EPI and PrE, initially distributed in a salt- and-pepper pattern. Near the onset of implantation, the PrE forms an epithelium separating the EPI and blastocoel cavity^9, 11, 13, 89^. **(B)** Representative blastocysts of good morphology (GM) and poor morphology (PM) by standard assessment^20^. A GM blastocyst comprises many cells forming a tightly-packed, well-demarcated ICM (indicated by arrowhead) and a cohesive TE epithelium. In contrast, PM blastocysts contain fewer, more loosely packed cells in both the ICM (indicated by arrowhead) and TE, often with fragmentation. **(C)** Experimental set-up. Embryos were cultured to blastocyst, graded per standard morphological criteria^20^, and vitrified to allow for classification as euploid by PGT-A (see Methods). After being thawed and allowed to expand, blastocysts were separated by laser dissection along the abembryonic(AB)-embryonic(EM) axis, and subjected to RNA sequencing. **(D-E)** A developmental trajectory for single blastocyst cells (D3-7) is plotted in pseudotime space. Both GM and PM **(D)** abembryonic and **(E)** embryonic samples lie on the developmental trajectory most consistent with D6 samples. **(F-G)** Gene set enrichment analysis (GSEA) for genes most significantly upregulated (ranked by signed p-value, see Methods) in GM abembryonic **(F)** and embryonic compartments **(G)**. These gene sets represent enriched biological gene ontologies (adjusted p-value < 0.05) for genes at the top of the ranked gene list for each embryo compartment, corresponding to genes most significantly upregulated in GM embryos.

It has been known for decades in the IVF field that blastocyst morphology correlates with implantation success—specifically, that blastocysts of “good” morphology exhibit a high potential for implantation and ongoing pregnancy, on average, and blastocysts with “poor” morphology exhibit a lower potential for success, on average^7, 19^. In fact, morphological assessment remains the primary means of selecting an embryo with the highest potential for successful implantation for transfer in IVF clinics worldwide. Decades of research have established a standardized morphologic grading algorithm based on assessment by microscopy^20^. Components of this algorithm include determination of embryo stage and expansion status (i.e., size of blastocoel cavity in relation to volume of the embryo) and assignment of a grade based on the number and morphologic appearance (i.e., cell organization and fragmentation) of cells within the ICM and TE. These grades together generate a composite grade and classification of good, fair or poor morphology^20^ (Fig 1B, S1A) to inform the choice of embryo for transfer, with good morphology embryos being preferentially transferred when available. Morphology, however, is not a perfect predictor, and blastocysts of “poor” morphology can successfully implant, albeit at significantly lower rates. Despite this long known and well-established difference in implantation potential based on microscopic morphological assessment, what distinguishes blastocysts of good and poor morphology at the molecular level remains essentially unknown. Therefore, identification of the molecular differences between these two groups of embryos provides a valuable opportunity to advance our understanding of the factors and pathways that influence implantation success in humans. Towards this end, we analyzed human blastocysts of divergent morphological grades and implantation potential to identify the subset of factors and cell-cell interactions specifically enriched in blastocysts with increased implantation potential.

## Results

### Good and poor morphology human blastocysts differ most significantly in rates of implantation

To validate the relationship between morphology and implantation potential in our clinic, we performed a retrospective analysis of pregnancy outcomes for 888 blastocysts following single- embryo transfer in IVF cycles over an 8-year period. Importantly, only blastocysts predicted to be euploid by standard preimplantation genetic testing in the clinic were included to minimize the known effects of aneuploidy on embryo morphology and implantation potential. Consistent with the known increased potential of good morphology (GM) blastocysts, transfer of GM blastocysts led to higher rates of live birth relative to poor morphology (PM) blastocysts (65.4% for GM vs. 30.0% for PM, p<0.001) (Fig S1B).

To determine the stage at which poor morphology embryos most commonly fail, we compared measurable causes of pregnancy failure in the IVF setting, including negative pregnancy test (defined as no detectable β-hCG production 11 days after embryo transfer), biochemical pregnancy (defined as transient β-hCG detection), spontaneous abortion (defined as fetal loss at 6- 20 weeks), and loss at stages greater than 20 weeks. We found that transfer of PM blastocysts resulted in a negative pregnancy test in patients at more than 3-fold higher rates than transfer of GM blastocysts (48.6% for PM vs. 15.1% for GM, p<0.001) (Fig S1B). As β-hCG production by invading TE cells is the first detectable sign of implantation in the clinical setting, these findings demonstrated a lower potential of PM embryos, as a group, to initiate or sustain the earliest stages of implantation^2, 21^. In contrast, the rates of both biochemical pregnancy (10.9% GM vs. 15.7% PM; p=0.125) and spontaneous abortion (8.6% GM vs. 5.7% PM; p=0.789) were similar between GM and PM blastocyst transfers (Fig S1B). Loss at stages later than 20 weeks was negligible in both groups. Together, these data demonstrate that the primary difference between the developmental potential of GM and PM embryos is in their ability to implant.

We next probed the relative contributions of the TE and ICM to implantation by reanalyzing these data by morphological grades assigned independently to the ICM and TE (A, B or C)^20^ (Fig S1A). Embryos with a C-graded ICM resulted in significantly higher rates of failed implantation (negative β-hCG for 59%) relative to embryos with a C-graded TE (negative β-hCG for 38%) (failed implantation p=0.039) (Fig S1C). Because GM embryos are preferentially transferred when available, our analysis is limited by the lower number of PM embryos transferred (n=135, C-TE; n=27, C-ICM). Nevertheless, these data suggest that the developmental state and morphology of the ICM—which does not directly contact the maternal endometrium—more commonly determines implantation success.

### Gene expression profiles enriched in human blastocysts of high implantation potential

To explore the molecular mechanisms underlying the significantly higher implantation potential of GM embryos, we selected two separate cohorts of vitrified supernumerary blastocysts of GM and PM donated for research by IVF patients in our clinic—one for transcriptomic analysis and a second for immunofluorescence analysis (totals: GM, n=9; PM, n=11; Data S1). All embryos in the final selection were fully expanded blastocysts at D5 at the time of vitrification (except for one D6 PM blastocyst in the transcriptome group, which was at the same developmental stage by morphologic analysis). The blastocysts were thawed and cultured for an additional eight hours on average to allow for full expansion, confirmation of viability, and a second assessment of morphological grade and matched developmental stage by a trained embryologist (see Methods). Thus, at the time of analysis, these embryos represented the developmental stage immediately following standard embryo transfer and, therefore, the molecular state as close to initiation of implantation as possible. In addition, to minimize effects due to chromosomal abnormalities, all selected blastocysts were reported as euploid by standard preimplantation genetic testing for aneuploidy (PGT-A) approaches in the IVF clinic before vitrification (the same approaches used prior to embryo transfer in Figs S1B-C, see Methods). Although this is the maximum extent of evaluation possible in the clinic and the current standard of care, a single biopsy cannot evaluate mosaicism in the remainder of the embryo. Therefore, we also screened embryos for detectable chromosome imbalances following RNA sequencing (see Methods and results below). Because blastocysts of good morphology and of normal PGT-A chromosomal screening currently remains the highest metric to enrich for embryos with the highest implantation potential, we reasoned that GM blastocysts, relative to PM blastocysts, would also be enriched for factors and pathways associated with successful implantation.

In preparation for RNA sequencing, each blastocyst was separated across its abembryonic- embryonic axis into two major compartments: (i) the mural TE [mTE, lies opposite the ICM], which will be referred to as the abembryonic (AB) compartment, and (ii) the ICM/pTE (comprised of cells of the EPI, PrE, and polar TE [pTE, overlies the ICM]), which will be referred to collectively as the embryonic (EM) compartment (Fig 1C). Our rationale for bisection of blastocysts via this method was two-fold: (i) to maximize proportional representation of all cells present in the embryo to facilitate comparison between GM and PM samples, and (ii) to allow for deeper sequencing of individual compartments than possible with single cell methods. With respect to (i), since multiple single cell human blastocysts datasets are already available, and because dissociation of blastocysts for single cell methods routinely captures ∼10%^15, 16, 22, 23^ of total cells per embryo, we concluded that assessment of total lineage contribution per embryo was most feasible via a bisection approach. We sequenced matched AB-EM compartments for a cohort of 14 stage-matched, euploid embryos with disparate morphological grades (6 GM and 8 PM). Only blastocysts for which a clear ICM could be identified and a clear bisection feasible were selected for sequencing. We also validated proper separation of the compartments bioinformatically, demonstrating that embryonic and abembryonic samples from both GM and PM embryos were enriched and depleted for expected lineage markers^16^ (Fig S2A-D, Data S2) and distinguishable by principal component analysis (PCA) (Fig S2E). Abembryonic samples from GM and PM blastocysts clustered more closely together than embryonic samples, indicating that gene expression in the embryonic compartment was more variable between GM and PM blastocysts relative to the abembryonic compartment (Fig S2E). With respect to developmental progression, GM and PM samples most closely matched the distribution of D6 blastocysts from published profiles of late-stage blastocysts, confirming that both groups were matched at similar stages of late blastocyst development as predicted by microscopic stage determination^16^ (Fig 1D-E). To validate the PGT-A results and assess the remainder of the embryo for mosaicism, we examined chromosome-wide expression imbalances in our RNA sequencing data^24^. Although the sensitivity to identify small numbers of aneuploid cells in embryo samples has not been demonstrated directly via this approach, we did identify one potential mosaic aneuploidy in a single GM abembryonic sample, which was retained in our final analysis as the proportion of mosaicism was <30%, not detected in the matched embryonic sample, and therefore, predicted to not affect developmental potential^25^ (See Methods; Fig S3).

We next compared the transcriptomes of abembryonic and embryonic compartments of GM and PM blastocysts to define molecular factors enriched in embryos of increased implantation potential (Data S3). In the abembryonic compartment, gene expression in GM versus PM blastocysts was highly correlated overall (R^2^ = 0.98), as was expression of a panel of TE-specific genes (n=98)^16^ (R^2^=0.96; p=0.21) (Fig 2A). Genes most significantly upregulated in GM relative to PM abembryonic samples were enriched for regulators of metabolism, translation, and cell cycle progression (Fig 1F, S4A, Data S3), highlighting a requirement for regulation of metabolic rates to support protein synthesis and proliferation—the most energy-intensive processes in the cell. Previous studies have demonstrated a correlation both between amino acid utilization and embryo developmental competence^26^ and between metabolic rates in the blastocyst and live birth rates following embryo transfer^27^. Enriched categories related to energy production included genes involved in ‘oxidative phosphorylation’ (e.g., *ATP5MC1, ATP5MC2,* and *NDUFS8*), ‘cellular respiration’ (*ADSL, MDH2,* and *COQ9*), ‘electron transport chain’ (*GLRX, CYB5A,* and *AKR1A1*), and ‘amino acid metabolism’ (*GOT1, PCBD1, BCAT2, and IVD*). Also overrepresented in highly ranked genes were gene sets involved in protein synthesis, including ‘translation’ (*MRPL52, SECISBP2, MRPS27*, and *MRPL30*) and ‘rRNA and tRNA metabolism’ (*TSEN2, DARS2, UTP6,* and *POP5*), and gene sets involved in replication and nucleic acid production, including ‘regulation of cell cycle G2/M transition’ (*CDK4*, *TUBG1*, and *HAUS1*) and ‘purine metabolism’ (*ADSL, IMPDH2, MOCOS,* and *PAICS*). These findings suggest that enhanced metabolism and energy production in the mTE is associated with increased implantation potential.

**Fig. 2.**
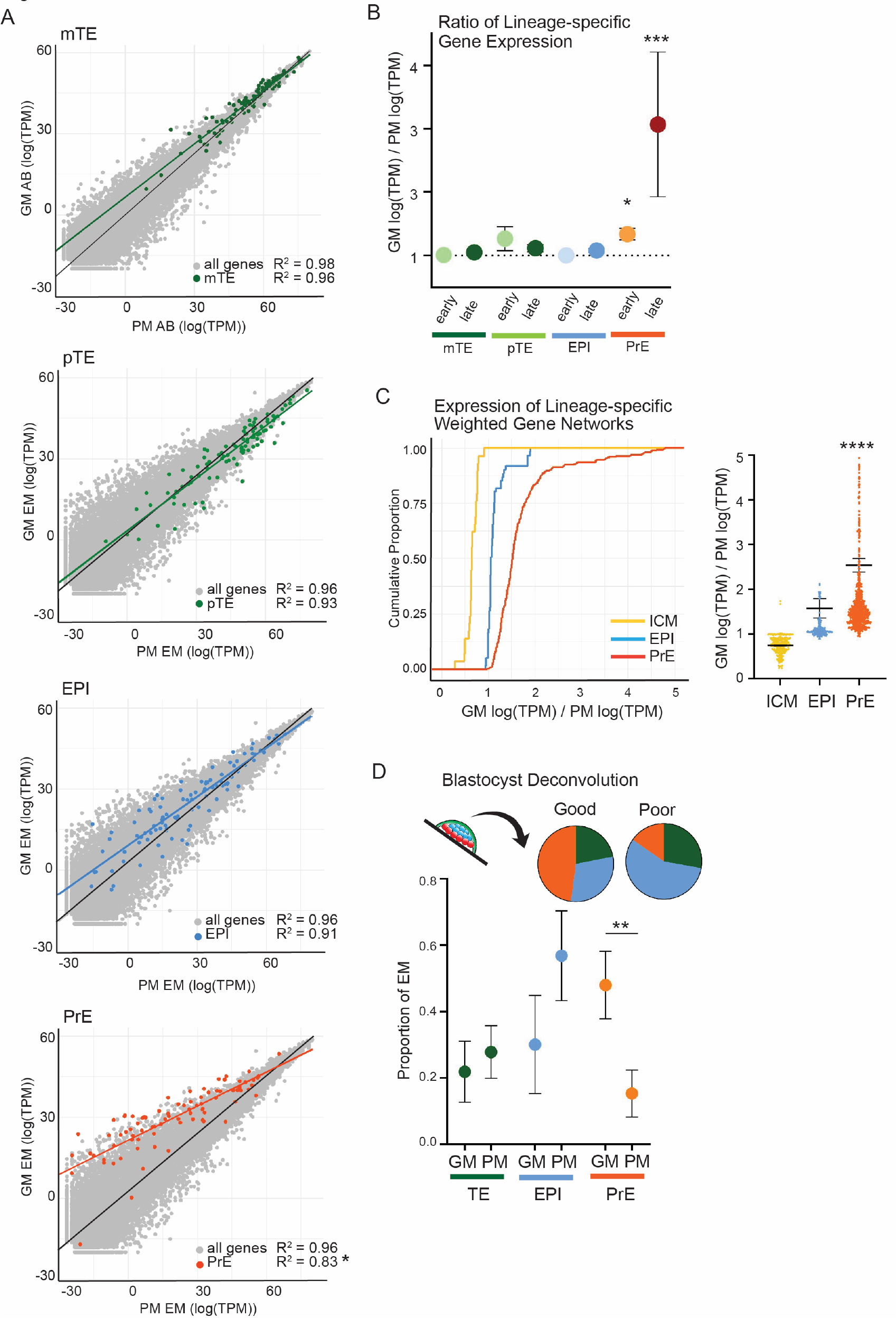
PM blastocysts demonstrate reduced expression of genes marking PrE development. **(A)** Scatter plots of the summed log(TPM) for genes expressed in the embryonic and abembryonic compartments of GM and PM embryos (grey). Lineage-specific genes^16^ for mTE (p=0.21, n=98), pTE (p=0.25, n=98), EPI (p=0.16, n=99), and PrE (p=0.04, n=82), are overlaid in the color indicated. *p<0.05, Student’s T-test. **(B)** Ratio of the log(TPM) for lineage-specific genes in GM versus PM blastocysts. Mean and standard error indicated. Mean ratios of established early and late developmental lineage markers^16^ are shown: mTE.early 1.01 +/- 0.05 (P=0.40), mTE.late 1.05 +/- 0.08 (p=0.18), pTE.early 1.27 +/- 0.63 (p=0.34), pTE.late 1.12 +/- 0.31 (p=0.30), EPI.early 1.00 +/- 0.13 (p=0.46), EPI.late 1.08 +/- 0.24 (0.41), PrE.early 1.33 +/- 0.32 (p=0.01), PrE.late 3.07 +/- 0.32 (p=0.0003). *p<0.05, *** p<0.001, Student’s T-test. n=14 total blastocysts. **(C)** Weighted gene network cluster analysis (WGNCA) was used to define lineage-specific gene expression networks for the pre-differentiated ICM and the differentiated EPI and PrE^14^. The ratio of gene nodes in GM versus PM embryonic samples is plotted as a cumulative distribution plot (left) and scatter plot (right). As the expression of a gene set in PM embryos decreases, the GM/PM ratio increases, and the cumulative frequency is shifted to the right, as is seen for the PrE network. **** Bonferroni- adjusted p-value = 9.8e-39 for GM versus PM PrE, p=0.26 ICM, p=1 EPI, Student’s T-test. **(D)** The proportion of each lineage contributing to GM and PM embryonic samples was predicted using a machine learning approach^31^: GM embryonic samples 22% pTE, 30% EPI, and 48% PrE (R=0.63); PM embryonic samples 28% pTE, 57% EPI, and 15% PrE (R=0.76). TE p=0.31, EPI p=0.11, PrE p=0.009. **p<0.01, Student’s T-test.

In the embryonic compartment, gene expression in GM and PM blastocysts was also highly correlated overall (R^2^=0.96) (Fig 2A). However, examination of panels of markers for each lineage independently revealed that PrE-specific genes (n=82)^16^ were expressed overall at much lower levels in PM relative to GM blastocysts (R^2^=0.83; p=0.04) (Fig 2A). This was in contrast to panels of both pTE-specific genes (n=98)^16^ (R^2^=0.93; p=0.25) (Fig 2A) and EPI-specific genes (n=99)^16^ (R^2^=0.91; p=0.16) (Fig 2A), which were expressed at similar levels in GM versus PM blastocysts. Markers of PrE development enriched in GM blastocysts included the highly conserved PrE marker *GATA4* and other established markers *(e.g., LINC00261, COL4A2, FN1, NID2, FLRT3,* and *PITX2).* Many of these genes have roles in extracellular matrix (ECM) formation, consistent with published findings linking ECM formation and cell adhesion with PrE development in model organisms^28–31^. In fact, the top gene sets enriched in GM relative to PM embryonic samples overall included ‘cell-cell adhesion’ (*CDH11, NECTIN3, DSCAML1,* and *AMIGO2*), pointing to a strong dependence upon cell-cell interactions and extracellular signaling within the embryonic compartment (Fig 1G, S4B). Also enriched in GM embryonic samples was ‘regulation of MAPK cascade’ (*GATA4, CAV1, FGB, ITGA1, MAP3K3, NRG1,* and *LBH*), a pathway critical for PrE development in mouse^30^, although its role in human PrE development remains unclear. Other gene sets among the those most enriched in GM embryonic samples included ‘carboxylic acid metabolism’, encompassing many factors involved in lipid metabolism (*HSD17B4, PPARG*, and *CRAT*) and amino acid metabolism (*P4HA1, HIBCH, HGD*, *SHMT2,* and *GPT2*), among other metabolic pathways. A resource containing the complete list of genes and gene categories enriched in each compartment in GM relative to PM embryos can be found in Data S3. Together, these data highlight the most significant genes and gene categories enriched in human blastocysts of good morphology and high implantation potential.

### Human blastocysts of reduced implantation potential demonstrate defects in primitive endoderm development

While our gene expression analyses of GM and PM blastocysts revealed a number of specific factors that distinguish embryos in these two groups, one of the major differences was reduced expression of a panel genes marking PrE development in PM embryos. To more closely examine differences in PrE-specific genes in the context of the other lineages in the blastocyst, we extended our analysis to look independently at established early and late markers of maturation for each lineage. Early markers are those detectable before the D5 blastocyst stage and indicative of lineage specification, while late markers only become detectable at D5 or later and are indicative of lineage maturation (see Methods)^16^. These analyses revealed that embryonic samples of PM blastocysts expressed lower levels of both early (p=0.02) and late (p=0.0003) PrE markers relative to embryonic GM blastocyst samples (Fig 2B). In contrast, GM and PM blastocysts exhibited similar expression of early and late markers for mTE, pTE, and EPI. Repeating these analyses using an additional published set of markers^14^ demonstrated a similar pattern, with a robust reduction in late PrE marker expression in PM embryonic samples (p=2.9e-06) and a more subtle but significant reduction in early PrE markers (p=0.03) in the absence of a significant difference in early or late EPI markers (Fig S5A). Because PrE development is closely associated with EPI development, we asked whether PM blastocysts might have EPI defects not apparent in our pooled analyses. Examining lineage marker expression within each embryo individually, we found that the reduction in both early and late PrE marker expression was consistent among PM blastocysts (Fig S5B). In contrast, expression of EPI markers was much more variable (Fig S5B). This variability was present in both GM and PM embryos and decreases in EPI marker expression were not specific to or consistent among PM blastocysts.

To examine the possibility of a preceding ICM defect in PM blastocysts, we evaluated gene expression in the ICM before and after EPI and PrE differentiation using a comprehensive gene co- expression network^14^. This study pooled blastocyst single-cell expression data from three large, published studies to define weighted gene networks of highly variable genes strongly linked to the early (pre-differentiated) ICM in contrast to the mature EPI and PrE lineages. Evaluating the most highly weighted gene nodes identified for each network (weight >0.1, n=307 early-ICM, n=222 EPI, n=917 PrE nodes) demonstrated that genes of the PrE network were expressed at significantly lower levels in PM relative to GM blastocysts, whereas genes of the EPI and early ICM were detected at similar levels (Fig 2C). These data revealed no consistent molecular differences in early ICM development in GM and PM blastocysts.

Because culture conditions and subtleties in embryo grading can vary between IVF clinics, we next asked whether the molecular differences we found in GM and PM embryos from our clinic are seen more broadly by examining a larger group of human embryos. Towards this end, we identified and analyzed published transcriptome data from a distinct cohort of embryos with reported morphology assessments from a different clinic^32^, comprising 20 whole blastocysts of GM (defined as A or B grade) and PM (defined as C grade) (GM, n=7; PM, n=13; Data S1). Despite the small number of cells represented by the PrE within whole blastocysts, the largest defect in PM relative to GM blastocysts was again in the expression of genes within the PrE network, with smaller EPI network defects (Fig S6A). Together, these analyses—in two distinct embryo cohorts from different clinics—demonstrate that the greatest and most consistent differences at the molecular level between GM and PM blastocysts is decreased expression of broad panels of genes marking development and maturation of the PrE.

The most likely explanation for the observed reduction in the expression of broad panels of PrE-centric genes in PM blastocysts, is a decreased number of PrE cells. To probe this possibility, we used both a computational and an experimental approach. First, we leveraged a machine learning deconvolution algorithm^31^, using a published single-cell blastocyst dataset in which cell types had already been defined as the training dataset^16^. This established ‘gene signatures’ for each lineage (pTE, EPI, and PrE). We then applied these signatures to quantify the relative proportion of cells of each lineage that contribute to each embryonic sample as a whole. This analysis revealed that GM samples contain a greater proportion of PrE cells, comprising 48% of GM embryonic samples (22% pTE, 30% EPI, 48% PrE) compared to only a 15% PrE contribution for PM embryonic samples (28% pTE, 57% EPI, 15% PrE) (Fig 2D). We also conducted this analysis using the second dataset of whole blastocysts^32^ (with a training dataset of TE, EPI, and PrE^16^), which revealed a similar reduction in the predicted PrE cell contribution in PM blastocysts, with PrE contributing 37% in GM blastocysts (35% TE, 28% EPI, 37% PrE) compared to only 13% in PM blastocysts (62% TE, 25% EPI, 13% PrE) (Fig S6B).

Second, we asked whether PM blastocysts contain a lower number of PrE cells experimentally at the protein level in a third cohort of blastocysts (GM, n=3; PM, n=3; Data S1). To do this, we performed immunofluorescence using lineage markers that are well-established and define each of the three lineages (CDX2 for TE, NANOG for EPI, GATA4 for PrE) ^9, 11, 33^. To ensure the relative proportion of cells of each lineage was represented, each embryo was co-stained for markers to all three lineages, followed by imaging and cell counting to quantify the number and relative representation of TE, EPI, and PrE cells within the embryo. Consistent with our previous analyses, we found that PM blastocysts contain almost 5-fold fewer PrE cells than GM blastocysts (GM 8.3 +/- 1.9 cells (6.9% of total cells per embryo), PM 1.7 +/- 0.5 cells (1.7%), 4.8-fold difference, p=0.004), with no significant difference detected in the number of EPI (GM 11 +/- 3 (9.1%), PM 8.7 +/- 2 (8.6%), 1.2-fold difference, p=0.215) or TE cells (GM 103 +/- 22 (85.4%), PM 93 +/- 20 (92.7%), 1.1-fold difference, p=0.334), or in the total number of cells (GM 121 +/- 21 cells, PM 101 +/- 22, 1.2-fold difference, p=0.201) (Fig 3A-C, S7A-B). Notably, we observed a trend towards decreased total cell number in PM blastocysts, as expected^20^. However, this was not a statistically significant finding in our cohort, likely due to the fact that we selected only PM embryos that were able to successfully develop to the late blastocyst stage, as well as the low number of stage-matched embryos available for analysis. Nevertheless, these data provided independent validation of our transcriptome analyses at the protein level and confirmed the presence of a significantly lower number of cells representing the PrE lineage in PM blastocysts.

**Fig. 3.**
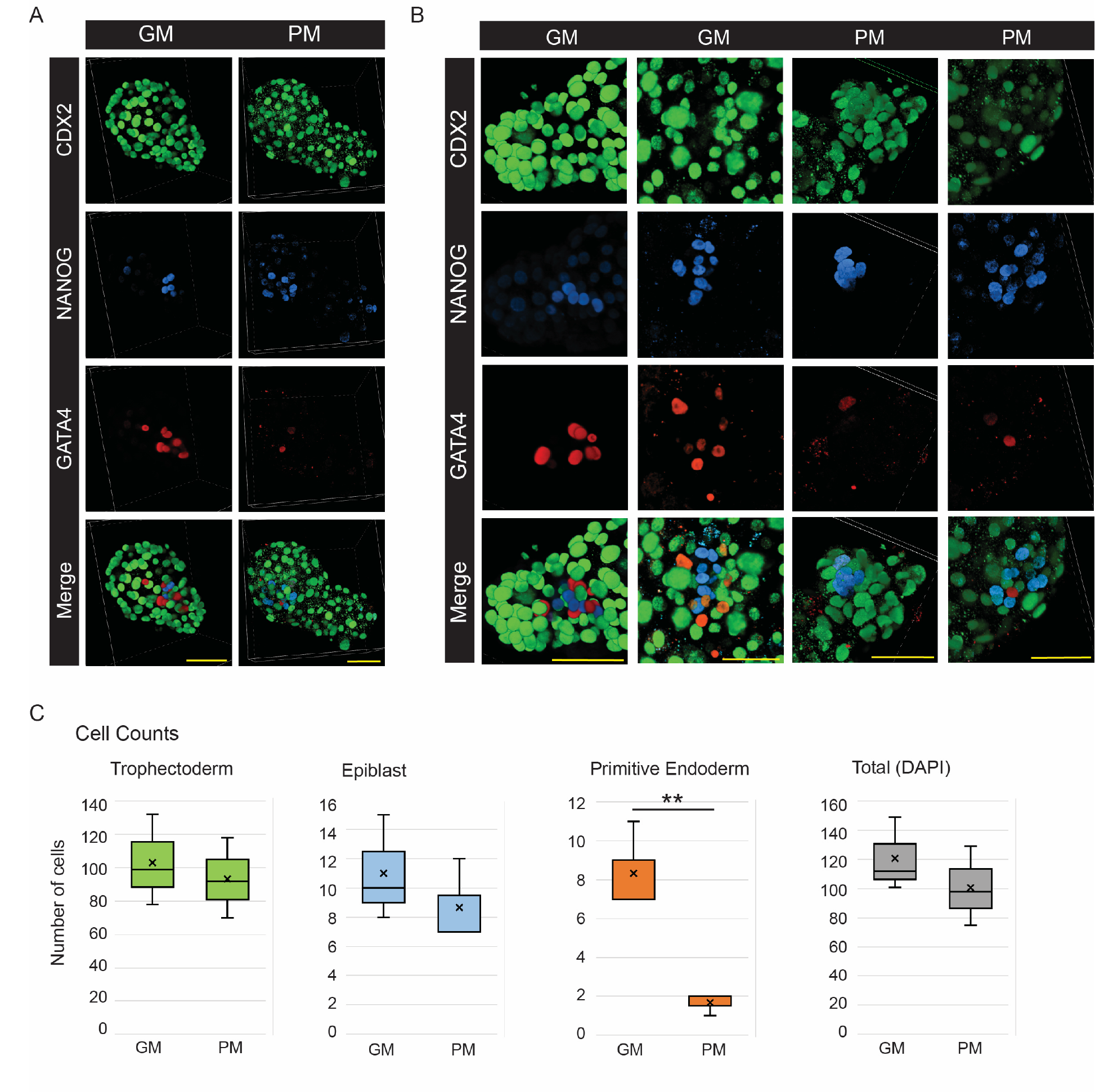
PM blastocysts are composed of a reduced number of PrE cells. (A) 3D volume projections of representative GM and PM blastocysts stained for markers of each lineage: CDX2 (TE), NANOG (EPI), GATA4 (PrE). Yellow scale bars = 50µm along 3D grid. **(B)** Zoomed 3D volume projections to allow better visualization of the ICM. Yellow scale bars = 50µm along 3D grid. **(C)** Nuclear (cell) counts for each lineage and total cell number per embryo. Median counts are indicated by the horizontal line within each bar, whereas mean counts are indicated by an “x”. (n=3 GM, 3 PM embryos) (mean +/- standard deviation: TE GM 103 +/- 22, PM 93 +/- 20, p=0.33; EPI GM 11 +/- 3, PM 8.7 +/- 2, p=0.22; PrE GM 8.3 +/- 1.9, PM 1.7 +/- 0.5, p=0.004; DAPI GM 121 +/- 21, PM 101 +/- 22, p=0.20). ** p<0.005, Student’s T-test.

In sum, we present multiple lines of evidence that point to common defects in PrE development at the time of embryo transfer among blastocysts of poor morphology and reduced implantation potential. First, our transcriptome analyses using large panels of early and late lineage- specific markers, as well as more comprehensive weighted gene networks in two distinct cohorts of blastocysts from different IVF clinics, showed consistent defects in expression of PrE-centric markers in PM blastocysts relative to GM blastocysts. Second, lineage-specific gene signatures identified via a machine learning approach provided a separate and unbiased approach and revealed that PM samples contain a smaller proportion of cells representing the PrE, with similar PrE deficits identified in both embryo cohorts. Third, immunofluorescence analyses examining the relative contribution of each lineage confirmed a reduced number of PrE cells in PM blastocysts in a third, independent cohort of embryos. That defects in PrE development were the most significant and consistent difference detected between GM and PM blastocysts suggests that the developmental state of the embryonic compartment, and of the PrE, in particular, is strongly coupled with both clinical morphology and implantation potential in the human blastocyst.

### Predicted cell-cell communication patterns identify signals enriched in human blastocysts of high implantation potential

Communication between and among cells of each lineage in the blastocyst is essential for successful embryo growth and implantation. These interactions include paracrine, autocrine, and juxtacrine signaling pathways, as well as direct cell-to-cell structural connections, and have been shown to mediate cell fate decisions^34–38^. In addition, while less studied, signaling factors are also likely transported across the blastocoel fluid alone or within extracellular vesicles from cells of the embryonic to the abembryonic compartment and vice versa to regulate cell fate and other cell processes^39, 40^. Investigation of these cell signaling pathways in human embryos is challenging on multiple fronts, including the technical and ethical challenges involved in perturbing gene expression to test the role of individual signaling factors and the large number of embryos required to perform these functional experiments. Most fundamentally, however, these challenges are exacerbated by the fact that the repertoire of cell-cell interactions within the blastocyst upon implantation—and the subset of these cell signals required for successful implantation—remains largely unknown.

To begin to address these challenges, we expanded our transcriptome analyses to delineate the network of cell-cell signals expected in GM embryos of high implantation potential. Further, by comparing cell-cell signals in GM and PM embryos, we identify signals enriched in embryos of high implantation potential and commonly absent in embryos less likely to implant. To do this, we leveraged a published repository of ligand-receptor interactions within a statistical framework^41^. By examining the frequency and expression of genes encoding ligand and receptor pairs, this algorithm predicts the likelihood that a signaling pair is active (i.e., that it is co-expressed at a level higher than expected by pure chance). In total, our analysis uncovered 213 predicted interactions within and between compartments of GM blastocysts (p<0.05) (Fig 4A, S8A and Data S4). Of those, 67% involved the regulation of cell and growth factor signaling, angiogenesis, immune regulation, and cell migration, with the majority of these signals originating from (i.e., the ligand was expressed in) the embryonic compartment. The remaining 33% of the interactions predicted in GM blastocysts involved ECM or cell-cell adhesion interactions. These signals also originated almost exclusively within the embryonic compartment, highlighting an abundance of structural interactions within the ICM in the late blastocyst. In total, the proportion of interactions originating from the embryonic compartment (74%) far outweighed those originating from the abembryonic compartment (26%), further reinforcing the importance of the ICM as a signaling hub within the blastocyst.

**Fig. 4.**
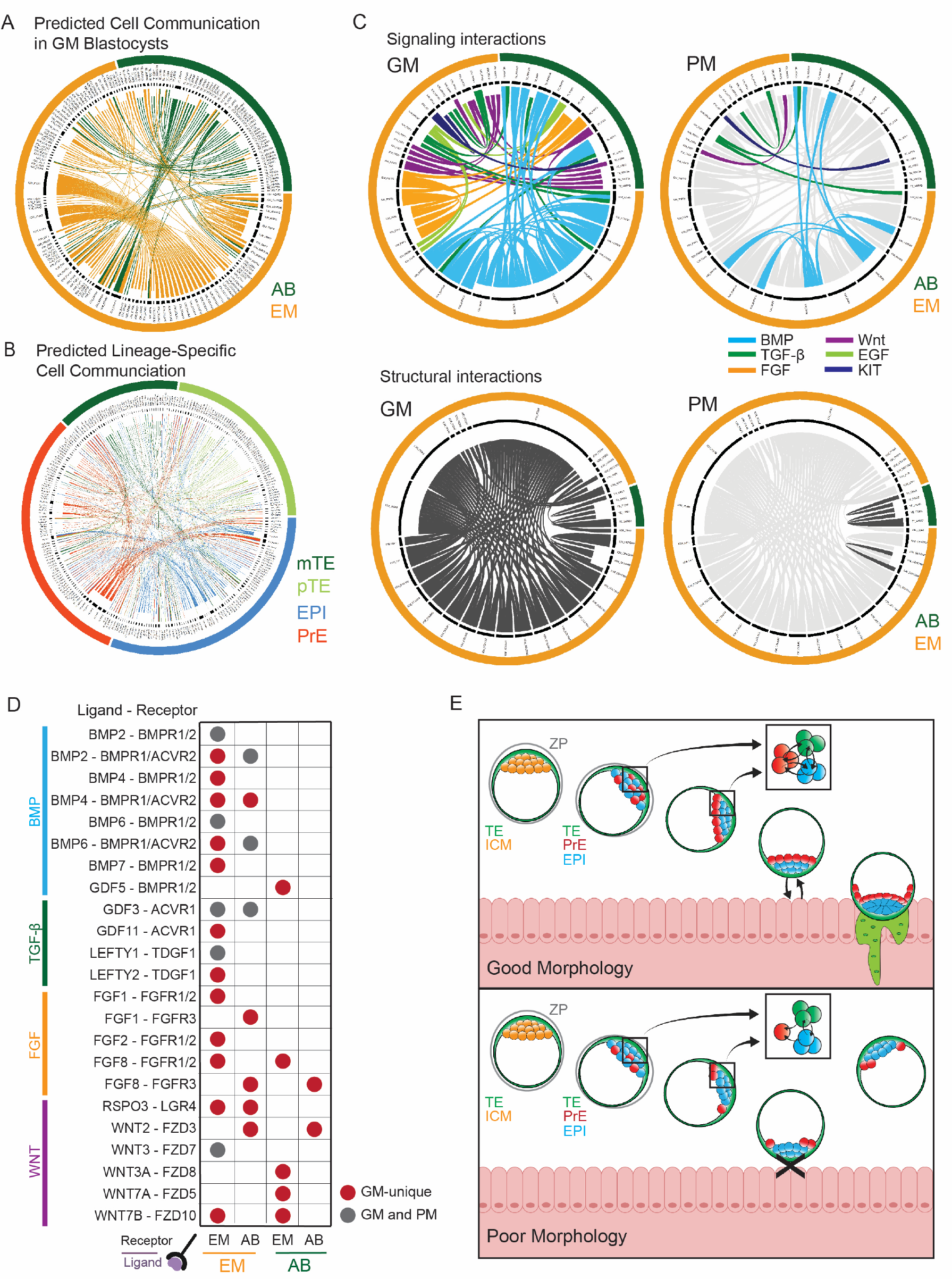
Cell-cell communication networks predicted in good morphology human blastocysts. **(A)** Cell-cell interactions predicted within GM embryos (p<0.05). Each line represents one ligand- receptor interaction, with interactions originating from the embryonic (EM) compartment (i.e., ligand expressed) in yellow and interactions originating from the abembryonic (AB) compartment in green, and the indented portion of the line marking the ligand. **(B)** Lineage-specific interactions via analysis of single blastocyst cells^16^. As in (A), interaction lines are directional, with the indented portion of the line marking the ligand. **(C)** A subset of cell-cell interactions predicted in GM (left) versus PM (right) embryos, revealing significant differences in signaling pathways and structural interactions. **(D)** Cell-cell signaling interactions predicted in GM embryos, including prominent members of the BMP, TGF-β, FGF, and WNT pathways. Ligand-receptor pairs predicted only in GM embryos are shown in red, and those also predicted in PM embryos are shown in grey. **(E)** Working model for the role of PrE in human blastocyst implantation. Here, we discover defects in PrE development and the PrE-derived ECM as the most significant and common difference between PM and GM blastocysts. Since PM and GM blastocysts are known to differ in their rates of implantation, our data demonstrate that the state of the PrE at the time of implantation is highly associated with implantation success. Because the PrE does not directly interact with the maternal endometrium during implantation, these data point to a model in which PrE-derived signals play important early roles in the acquisition of pTE implantation competence and, ultimately, in successful embryo implantation in humans.

We also conducted this analysis on a published dataset of single blastocyst cells^16^, which allowed further delineation of cell-cell signaling interactions between and among cells of each lineage and demonstrated a similar distribution of signals compared to analysis of our samples. This analysis uncovered 501 predicted lineage-specific interactions (p<0.05) (Fig 4B, S9, Data S4). Highlighting again the abundant signaling within the embryonic compartment, the majority (∼75%) of predicted ligand-receptor interactions within the blastocyst originated from the EPI and PrE, with a similar number of signals predicted to arise from each (EPI, n=184 (37%) and PrE, n=192 (38%)). In contrast, fewer ligand-receptor interactions were predicted to originate from cells of the TE, where 16% (n=78) of predicted interactions originated from the pTE and 10% (n=48) from mTE. A resource containing a complete map of lineage-specific interactions predicted in human blastocysts^16^ is provided to focus future studies examining specific signaling pathways in human embryos (Data S4).

Next, to identify signals specifically enriched in blastocysts of increased implantation potential, we compared ligand-receptor interactions predicted in GM versus PM blastocysts. We found that 80% (n=171) of interactions predicted in GM blastocysts were not predicted in PM blastocysts. The majority (79%) of these ‘GM-unique’ signals originated from the embryonic compartment (Fig 4C-D, Data S4), further supporting the idea that the defects within PM blastocysts arise predominantly within the embryonic compartment, as suggested by analyses of our clinical transfer data (see Fig S1C). Signaling interactions specifically enriched in GM blastocysts included pathways well-studied in embryos, such as BMP, FGF, TGF-β, and WNT, and we identified specific candidate factors within these pathways to focus future studies. These data also identified pathways less studied to date, including KIT, EGF, and Ephrin (Fig 4C-D), providing new directions for investigation as well.

Given that the greatest difference between GM and PM blastocysts was in the state of the PrE, we would expect signals most highly enriched in the embryonic compartment of GM blastocysts to include those important for PrE development—a process that remains poorly understood in humans. Consistent with this idea, we found a number of pathways implicated in PrE development, primarily in model organisms (PDGFA, BMP, FGF and WNT pathways), that were enriched in the embryonic compartment of GM blastocysts but not of PM blastocysts. These data validated our approach and pointed to specific factors in these pathways as candidates with important roles in humans (Fig 4C-D). For example, PDGF signaling regulates PrE maintenance and expansion in mice, with *Pdgfra-*null blastocysts exhibiting a reduced number of PrE cells and delayed implantation^42^. Consistent with a similar role for PDGF signaling in humans, multiple PDGF interactions were predicted in the GM embryonic compartment but none were detected in PM samples. Similarly, WNT signaling is known to play a key role in PrE development in primates^43–45^. The WNT family members *WNT7B* and *RSPO3* were uniquely enriched in GM samples (Fig 4C-D), suggesting a potential role for these specific factors in PrE development in humans.

In addition to factors driving PrE development, the embryonic compartment of GM blastocysts would also be expected to be enriched for signals that promote pTE implantation competence, the mechanisms for which also remain largely unknown in humans. Intriguingly, the FGF and BMP pathways have been shown to play a role in development of both the PrE and pTE in mice^46–48^, suggesting these lineages might develop in parallel and in response to shared signals from the EPI. Whether or not this is the case in humans remains unclear^13, 43^. With respect to the BMP pathway, our analyses identified the *BMP4* and *BMP7* ligands as ‘GM-unique’ (Fig 4C-D). In contrast, BMP signaling via other ligands (e.g., *BMP2* or *BMP6*) was similar in both GM and PM embryonic samples. In mice, knock-down of *Bmp4* and *Bmp7* leads to a dramatic reduction in the number of both TE and PrE cells^46^. These data suggest a similar role for *BMP4* and *BMP7* in human PrE—and potentially pTE—development, as well. Similarly, important roles for FGF signaling (predominantly FGF4) in PrE establishment and trophoblast stem cell maintenance have been defined in mouse^35, 38, 47–49^. Examining predicted interactions within the FGF pathway, *FGF2* and *FGF8* were uniquely predicted in GM embryonic samples (Fig 4C-D), suggesting roles for these specific FGF family members in PrE and/or pTE development in humans. Another group of prominent ‘GM-unique’ signals arising from the embryonic compartment included components of the TGFβ pathway known to be important for regulating pluripotency in EPI, including *GDF5, GDF11*, and *LEFTY2*^15, 50, 51^ (Fig 4C-D), raising the possibility that TGFβ signaling from the EPI might also contribute to processes affecting implantation potential.

Interactions that originated from the abembryonic compartment comprised the remaining 36 (21%) predicted ‘GM-unique’ interactions, with the large majority (81%; 29/36) from abembryonic to embryonic and only a small number predicted within the abembryonic compartment (19%; 7/36) (Fig 4C-D, S6A). As in the embryonic compartment, ‘GM-unique’ signals from the abembryonic compartment included pathways with well-established embryonic roles, namely FGF, TGF-β, and WNT. Indeed, WNT signaling was particularly prominent and unique to GM blastocysts, with specific factors predicted to signal from the abembryonic to the embryonic compartment (*WNT3A* and *WNT7A/B*) and others (*WNT2*) among cells within the abembryonic compartment itself. Future studies teasing apart the complex nature of WNT signaling dynamics within human TE will provide valuable insight into early placenta formation.

Strikingly, of the ECM and cell adhesion signals that comprise almost one-third of interactions predicted in GM blastocysts, the large majority (94%) were absent in PM blastocysts (Fig 4C). Within the embryonic compartment, the majority (52/66) of these structural interactions involved integrin-collagen connections (enriched integrin subunits: *ITGA1*, *ITGA2*, *ITGB1*; enriched collagens: *COL4A1*, *COL4A2*, *COL21A1*) (Fig 4C, S6B-C). In mice, the expression of ECM factors and PrE development are intimately linked^28, 52–54^. *Itgb1-*knockout mice demonstrate defects in PrE formation and peri-implantation lethality^28, 52–54^, and embryos lacking cell adhesion factors fail to develop and/or maintain the PrE^34, 55–59^. That PM blastocysts do not robustly express mRNAs encoding ECM and cell adhesion factors further supports a predominant PrE defect in PM blastocysts. This robust signature distinguishing GM and PM blastocysts points to PrE-associated structural factors as a prominent mark of blastocyst implantation competence.

To further validate signals predicted as enriched and depleted in GM and PM blastocysts, respectively, we asked whether the pattern of ‘GM-unique’ ligand-receptor pairs predicted in embryos from our fertility were similar to signals predicted within the separate cohort of blastocysts from a different clinic^32^. Analysis of the core functional categories (shown in Fig 4) demonstrated enriched expression of these genes, as a group, in GM versus PM blastocysts (Fig S10A-B). This enrichment was the most robust for genes with structural and signaling roles. We also examined ligand-encoding genes unique to each lineage^16^ (as shown in Fig S9). Plotting the ratio of lineage-specific ligand expression in GM versus PM embryonic compartments (our data) (Fig S10C), and in whole blastocyst samples (additional cohort)^32^ (Fig S10D), we again detected significantly reduced expression of PrE-associated ligands in PM blastocysts. These data further demonstrate consistent defects in PrE and the PrE-associated ECM among PM relative to GM human blastocysts. This resource containing a complete network of predicted cell-cell interactions enriched in GM blastocysts (representing drivers of normal development and implantation), as well as those unique to PM blastocysts (representing common developmental errors), is provided (Fig 4, S8-9 and Data S4).

## Discussion

Blastocyst morphological grade has been known for decades in the clinic to correlate with implantation success. However, what these morphological differences represent at the molecular level has remained unknown. This is an important question, as delineation of these molecular differences provides a rare and exciting opportunity to advance our understanding of mechanisms that underlie implantation competence and failure in humans. Here, we leverage this correlation between morphology and implantation potential, and through the direct comparison of good and poor morphology blastocysts, generate the first genome-wide list of factors and signaling pathways associated with implantation potential in humans. These datasets provide a rich resource for the field to direct future mechanistic studies in humans and in more genetically tractable model organisms aimed at uncovering the specific factors and pathways that underlie implantation competence of the human blastocyst.

While our analyses identified many molecular differences in gene expression and cell-cell signaling in GM versus PM blastocysts, the greatest difference between these embryos of disparate implantation potential was in the state of PrE development. Using multiple complementary approaches in three embryo cohorts from two different fertility clinics, our analyses consistently pointed to relative defects in factors and cell-cell signals derived from the embryonic compartment, particularly those associated with the PrE, among blastocysts with reduced implantation potential. At approximately Day 6 of development (see Fig 1D-E), these embryos represent the stage immediately following standard embryo transfer and as close to the initiation of implantation as possible. These defects in PrE development were detectable only at the molecular level and not discernable in otherwise stage-matched embryos during morphological grading by trained embryologists. In addition, they were largely limited to the PrE and detected in the absence of significant or consistent molecular defects in other blastocyst lineages. Finally, the defects in PrE development were also surprisingly consistent among the PM embryos examined, suggesting that this is a common defect in PM blastocysts and highly associated with human embryos of reduced implantation potential.

How might defects in PrE development in PM blastocysts arise? The etiology is likely multifactorial and may vary among embryos. For a subset of PM embryos, defects might lead to a delay in PrE development (as compared to GM blastocysts) in the late blastocyst, but with development still adequate and able to “catch up” in time for implantation. In support of this idea, almost one-third of PM blastocysts successfully implanted and resulted in live birth following embryo transfer (Figure 1C-D). This is direct evidence that a subset of PM blastocysts will develop a functional PrE with prolonged culture in an ideal environment (i.e., the uterus). Defects in other PM embryos might result in a more substantial PrE delay that is unable to “catch up” in time to implant successfully. In this case, if the state of PrE development is critical for the embryo to progress through implantation, any delay in the PrE might lead to a mismatch in the timing of implantation competence and endometrial receptivity and, therefore, to failed implantation. For the remainder of PM blastocysts, PrE defects might represent an irrecoverable failure of PrE differentiation, maintenance, or survival. These defects might be PrE-specific or a shared outcome downstream of varied ICM or EPI defects not robustly detected by our methods. In support of this possibility, while our analyses did not uncover consistent defects in EPI development, EPI marker expression was more variable than other lineages in both GM and PM embryos. This likely reflects both the variability inherent among human embryos and the variability of potential molecular defects that underlie poor morphology and poor PrE development. However, irrespective of the specific cause of the PrE defects, our findings demonstrate that the state of PrE development and implantation potential in the late blastocyst are strongly coupled and point to markers of the PrE, including the PrE-associated ECM, as a molecular signature of blastocyst implantation potential.

It is well-accepted that the blastocyst must be at the optimal developmental stage at the time of implantation, as this determines the confluence of factors and signals present to initiate and sustain interaction with the maternal endometrium. However, precisely what defines the optimal stage and what factors and pathways must be present to maximize implantation efficiency remains poorly understood. Our data suggest that the state of the PrE is a defining factor of that optimal stage. However, this raises the question of how development of the PrE, which does not directly interact with the maternal endometrium, is coupled with implantation potential. We suggest two possible models, both of which have interesting implications for our understanding of the process of implantation in humans. One possibility is that the PrE develops and matures in parallel with the TE in response to shared signals from the EPI, as has been demonstrated in mice^46–49^. Indeed, studies in stem cell-based human blastoids have shown that signals from the EPI directly contribute to pTE maturation and that this process is essential for attachment to endometrial cells *in vitro*^23^. While specific signals from the EPI that directly contribute to PrE differentiation in human embryos are less established^9, 33^, current data supports an important role for the EPI in PrE development^13, 14, 43^. Thus, in this model, the PrE would serve as a surrogate marker for implantation competence, without directly contributing to pTE differentiation or implantation success. Alternatively, given that differences in the PrE and the PrE-associated ECM were the greatest and most significant molecular differences in embryos with vastly different implantation potential— and much greater than differences in the pTE or EPI—it is tempting to speculate that the PrE itself plays an important role in the induction of pTE and implantation success. Consistent with this idea, the second and favored model is that competence of the pTE to mediate successful implantation requires integration of ICM-derived signals and that the PrE and/or the PrE-derived ECM critically contribute to these signals. A large body of work in mice has been devoted to the link between the PrE, embryo structural integrity, and resulting implantation competence^52, 55, 57, 58, 60, 61^. Accompanying lineage differentiation, the PrE begins to express extracellular matrix components that assist with PrE epithelium formation^59^. In addition, loss of any of a number of different ECM- or adhesion-related components in mouse blastocysts leads to a failure to develop, maintain, and/or epithelialize the PrE, with subsequent embryo lethality during peri-implantation development^52, 55, 57, 58, 60, 61^. Since these factors do not directly interact with the maternal endometrium, these findings are consistent with the idea that PrE- and/or PrE-associated ECM signals influence implantation via direct or indirect communication with the pTE. With large defects in PrE development and no significant differences detected in the early ICM or EPI, our findings support this body of published data and point to the second model, with a role for the PrE in mediating implantation (Figure 4E). Of course, these models are not mutually exclusive and components of each likely occur. Central to both models, however, is the association between the state and quality of the PrE at the time of implantation and implantation success.

The datasets provided here identifying factors and cell-cell interactions enriched and depleted in GM and PM blastocysts, respectively, pave the way for future mechanistic studies into the mechanisms that underlie successful implantation of the human embryo and the relationship between PrE development and implantation competence. The limited number of donated, chromosomally normal, stage-matched blastocysts of the highest and lowest quality available for these analyses is a limitation of this study. However, that multiple approaches in embryos from multiple clinics consistently identified similar molecular profiles for GM and PM embryos suggests that our analyses were able to identify the most significant and consistent differences in embryos of high and low implantation potential. Future studies comparing gene expression patterns and lineage marker analyses for GM and PM embryos across earlier stages of preimplantation development are needed to determine when preceding defects in PM embryos commonly arise and to identify contributing factors. Specifically, these experiments will probe whether defects in PrE in PM embryos observed here are due to errors in PrE specification, maturation, proliferation, and/or survival, as well as upstream mediators. In addition, as implantation occurs in the uterus, it is not currently possible to know at what point during the process of implantation that PM embryos most commonly fail. For instance, it is possible that PM embryos with deficient PrE are able to initiate implantation but are rapidly resorbed before implantation can be detected clinically. As protocols that allow *in vitro* culture of embryos beyond the blastocyst stage^10, 62^ become more efficient and feasible with a limited number of embryos, future studies that determine precisely when PM embryos most commonly fail and further elucidate the etiology of PrE defects—and other defects—in PM embryos will be important next steps. Finally, integration of our gene expression profiles and predicted cell-cell signaling networks from GM and PM embryos will identify candidate factors and pathways important for implantation to focus future mechanistic studies in genetically tractable models that allow functional testing. The recent development of human blastoid and other stem-cell based human embryo models provide exiting opportunities to rigorously test the role of these candidate factors, with the possibility of genetic manipulation in specific lineages and readout via *in vitro* implantation models^23^.

These data identifying factors and pathways enriched in blastocysts of the highest implantation potential have important implications for embryo culture in the clinic as well, with the potential to improve IVF outcomes. Specifically, the link between PrE development and implantation competence suggest that a better understanding of factors important for human PrE development could lead to optimization of media components to further promote or support PrE specification and maturation. Similarly, if it is found that the PrE itself provides signals important for implantation, media supplemented with PrE-derived factors might also improve implantation success. For the subset of PM embryos in which PrE development is delayed, embryo culture could be prolonged and transfer postponed to better match the timing of PrE development with the optimal window of endometrial receptivity. In support of this possibility, some studies have demonstrated that extended culture of PM embryos before transfer can increase implantation rates for a subset of PM blastocysts^63–65^. Whether the subset that does successfully implant after additional culture represents PM embryos with a PrE that is able to “catch up” during extended culture is an interesting question but will require the development of methods to assess the state of the PrE in live embryos before transfer to answer. Further, that extended culture does not rescue implantation for the majority of PM embryos in these studies suggests that defects in most PM embryos constitute more than a simple delay^63–65^.

That the PrE emerged from our studies as a potential marker for implantation potential also has implications for embryo selection. These findings raise the possibility that methods to assess the state of PrE development in live embryos before transfer might be a novel approach to improve selection of the embryo with the highest implantation potential for transfer. For example, detection of factors secreted by the PrE, present in spent culture media or in fluid collected from the blastocoel cavity, might allow for non-invasive assessment of PrE status before transfer. Alternatively, although we would predict a greater effect on pTE, it is possible that ICM- or PrE-derived effects on mTE could be detected via parallel transcriptome analysis of mTE biopsies commonly collected for genetic testing. Ultimately, prospective studies in which more comprehensive analyses of morphologic measures and/or gene expression profiles are performed prior to embryo transfer are needed to further delineate the confluence of factors distinguishing embryos of high and low implantation potential. Development and integration of methods such as these to improve embryo selection and increase implantation rates following transfer of a single embryo are critical to simultaneously reduce the risk of multiple pregnancy to both mother and fetus, and to improve success rates for a greater number of infertile couples. Together, our data begins to address these challenges, providing needed insight into common errors in development that underlie high rates of implantation failure for human embryos and the factors and mechanisms required for successful embryo implantation in humans.

## Materials and Methods

### Ethical Approval

Both the embryo research and retrospective analysis of pregnancy outcomes following single embryo transfer were performed under the approval and guidance of the Human Research Protections Program and Institutional Review Board at UC San Diego (IRB Project 191494).

Blastocysts used in this study were surplus, vitrified embryos from IVF treatment donated for research by patients in the IVF clinic following patient consent. Embryos were not generated for research purposes, and patients were not compensated for embryo donation.

### Clinical outcomes assessment

All embryos included in the retrospective pregnancy outcome analysis and selected for transcriptome analysis were graded based on morphological quality (Fig 1A) and trophectoderm biopsy by an experienced embryologist before vitrification according to standard procedures and protocols in the clinic. The ICM and the TE were graded independently (A, B, or C, where A is high, and C is low) ^66^. An embryo with either an ICM or a TE that was assigned a score of A was graded as “good morphology” whereas a score of C for either lineage resulted in a grade of “poor morphology” (Fig 1A, S1A-B). Trophectoderm biopsies (3-5 cells) were collected for preimplantation genetic testing for aneuploidy (PGT-A) in the course of standard care in the clinic before freezing by vitrification on day 5 or 6. PGT-A analysis was performed via either array comparative genomic hybridization (array CGH) or via next generation sequencing (NGS) by outside commercial laboratories (see below for further details). All single transfers of euploid blastocysts of good or poor morphology between 2012-2019 were included in the retrospective analysis. A few transfers in which 2 embryos were transferred were included if the outcome of each single embryo could be definitively determined by gender. After embryo transfer, clinical pregnancy outcome was tracked and recorded as *Negative Pregnancy Test* (human chorionic gonadotrophin (β-hCG) < 5 mIU/ml, 11 days after embryo transfer)*, Biochemical Pregnancy* (β- hCG > 5 mIU/ml (although typically <100) that does not rise normally, with no evidence of ectopic pregnancy)*, Spontaneous Abortion* (pregnancy documented by ultrasound at 6 weeks but lost before 20 weeks), or *Live Birth*.

### Blastocyst culture, morphological assessment and genetic screening

For the retrospective analysis of clinical pregnancy outcomes, all blastocysts were derived from ICSI-fertilized oocytes and cultured under standard conditions in the IVF Clinic (7%CO2/5%O2 in Vitrolife G1 media from Day 0 to 3 and in G2 media from Day 3 to 5-6 under mineral oil). All embryos underwent assisted hatching, which entails thinning of a single spot in the zona pellucida with a laser on Day 3 and is standard practice in the lab for all embryos that will undergo trophectoderm biopsy. The degree to which this affected the extent and timing of hatching is not precisely known but the treatment was the same for all embryos in both groups. Trophectoderm biopsies (3-5 cells) were collected for preimplantation genetic testing for aneuploidy (PGT-A) in the course of standard care in the clinic before freezing by vitrification on day 5 or 6 (all blastocysts were frozen on day 5 with the exception of one poor blastocyst frozen on day 6). PGT-A DNA analyses were performed via either array comparative genomic hybridization (ArrayCGH) prior to 2015, or via next generation sequencing (NGS) by Reprogenetics (prior to 2017) or Igenomix (2017 and later). Embryos with a mosaicism level less than 20% (Reprogenetics) or 30% (Igenomix) were reported to the clinic and patient as euploid and deemed appropriate for transfer ^25^. Embryos deemed euploid by these criteria were also considered euploid for the clinical embryo transfer outcomes’ analysis above and for the initial selection of embryos for our transcriptome analyses. While these screening tools are the current standard of care in the clinic, they are unable to detect all chromosome abnormalities, including some deletions, mutations, inversions, and translocations. A single TE biopsy is also unable to rule out a low level euploid/aneuploid mosaicism, for which the true incidence, the most accurate method of detection, and the actual clinical impact remains unknown^67^. We attempted to address this possibility to the degree possible with an additional mosaic aneuploidy assessment as described in more detail below. Prior to analysis, embryos were thawed according to standard protocols and cultured in G2 media (Vitrolife) and cultured for approximately 8 hours to confirm re-expansion and viability, at which time a final stage and morphological grade were assigned before laser dissection and freezing for analysis. Vitrification and thawing were conducted using the Cryotop method (Kitazato). An experienced embryologist separated the embryo along its anembryonic-embryonic axis by gently pulling the embryonic pole into the holding pipette and then using the laser across the base of the holding pipette to separate the embryonic pole from the anembryonic pole, which remained outside the pipette. Identifying and sequestering the embryonic pole in the holding pipette during dissection circumvented the challenge of identifying the ICM/embryonic pole once the integrity of the sphere is breeched and the blastocyst collapses. Each sample was individually washed in TE Wash Buffer (1mg/ml bovine serum albumin in 1XPBS), transferred to individual 0.2ml PCR tubes, and frozen immediately in liquid nitrogen. Importantly, only embryos in which the ICM could be clearly identified under the microscope were dissected and used for analysis, and marker analysis from RNA sequencing data was used to confirm dissection quality.

### RNA sequencing library preparation

In total, six GM and eight PM expanded human blastocysts were used for RNA sequencing. RNA isolation and cDNA synthesis were carried out using the SMARTer® Ultra® Low Input RNA Kit for Sequencing – v4 (Clontech, Cat.no. 634888). Libraries were generated from the resulting cDNA (0.2ng/ul per sample) using the Nextera XT DNA library preparation kit (Illumina, Cat.no. FC-131-1024) as previously described^68^. Indexed sequence libraries were pooled for multiplexing, normalized by MiSeq read number, and paired-end sequencing was performed on a HiSeq 4000.

### Analysis of differential gene expression

Approximately 18.1 million reads were generated per sample, and 70.1% of those were uniquely mapped via STAR (2.5.2a)^69, 70^ after trimming for adapter sequences via cutadapt (v1.10)^71^. Samtools^72^ was used to process sam files, as well as to sort and remove PCR duplicates of bam files. Counts for each gene were quantified using the Subread package FeatureCounts using the gene level quantification in paired-end mode (release 1.5.2)^73^, and annotated using the Ensembl GRCh38 genome. Genes were filtered such that genes without at least one sample with at least 10 raw reads were removed from the analysis. Pre-count data processing was performed on the Triton Shared Computing Cluster (TSCC)^74^. Count data was normalized using the Bioconductor package edgeR^75, 76^. The betweenLaneNormalization function of the Bioconductor package EDASeq^77^ was used to normalize the data using upper-quartile (UQ) normalization, and differential expression was calculated using DESeq2^78^. No bias in library complexity or percent of uniquely mapped reads per sample was observed between embryonic and abembryonic samples, or between GM and PM embryos. To account for multiple testing, genes with a Q-value less than 0.05 were considered differentially expressed. Heatmaps were constructed using the heatmap.2 function in ggplot2^79^. Principal components were calculated with the prcomp function. All box plots, scatter plots, PCA plots, and heatmaps were constructed using ggplot2. All these analyses and plot constructions were performed with RStudio, R (v3.4.0 and v3.5.2).

Supplementary tables of normalized read counts and calculated differential expression from Petropoulos et al.^16^, Blakeley et al.^15^ and Stirparo et al. ^14^ were used for the comparison of lineage-specific genes. Single-cell gene expression normalized TPM reads from Petropoulos et al.^16^ were used for lineage-specific marker identification, proportional composition analysis (CIBERSORT)^80^, generation of diffusion maps^81^, and cell compartment interaction analysis (CellPhoneDB)^41^. For these analyses, genes were filtered such that at least 50 single-cell samples must have a raw read count of at least 10.

### Gene Set Enrichment Analysis (GSEA)

Gene set enrichment analysis (GSEA)^82^ was performed with the Bioconductor package clusterProfiler^83^ using gene ontology biological process annotations, 1000 gene set permutations, and rank scores for differential expression calculated as the product of -log10(adjusted p-value) and the sign of the fold change, such that genes upregulated in GM embryos had positive scores and downregulated genes had negative scores. Significant gene sets were determined by a Benjamini-Hochberg adjusted p-value cutoff of <0.05 and then ranked by set size to reduce redundancy. The top 100 gene sets were visualized with the Cytoscape plug-in EnrichmentMap^84^ with default options and annotated with AutoAnnotate^85^ with manual renaming and layout adjustment for clarity.

### Mosaic Aneuploidy Assessment

To identify mosaic aneuploidies, we used a method that examines chromosome-wide expression imbalances, R package scploid ^24^. Briefly, this method calculates a “chromosome score” for each chromosome, measuring whether the expression level of genes in that chromosome differs substantially from other cells. Chromosome scores >1.2 are called as a gain, and scores <0.8 are called as a loss. This score is then converted into a p-value and used to detect significant deviations, providing evidence for the presence of an aneuploidy or mosaic aneuploidy.

### Pseudotime analysis

Pseudotime trajectories were constructed with the Monocle 2 package (v2.10.1)^86^. The ordering genes were identified as having high dispersion across cells (min_expression > 10; dispersion_empirical >= 1). For final plots, the top 500 overdispersed genes were used for ordering. The discriminative dimensionality reduction with trees (DDRTree) method was used to reduce data to two dimensions.

### Lineage marker analyses

A number of approaches, using marker genes from multiple studies, were utilized to assay each lineage of the human blastocyst. Marker genes as defined in Petropoulos et al. were obtained by combining Z-scores obtained from differential expression analysis of one lineage against each of the other two, and creating a ranked list of genes unique to the TE, EPI, and PrE ^16^. Petropoulos et al. also defined both “early” and “late” marker genes for each lineage, defined as being enriched in early (prior to D5) embryo samples, or late (after D5) embryo samples ^16^. These genes were used to construct lineage-specific developmental trajectories, diffusion maps, and their mean expression was additionally presented for our blastocyst samples. Marker genes as defined in Stirparo et al. were obtained via Weighted Gene Network Cluster Analysis ^87^ in which gene networks central to the pre-differentiated ICM, the EPI, and the PrE were compiled ^14^. These marker genes were used for cumulative distribution analysis of both our embryo samples and to further assess the ICM of an additional published cohort of blastocyst samples ^32^. We further used both “early” and “late” marker genes as defined by Stirparo et al. ^14^. Similarly to Petropoulos et al., early marker genes were expressed from the morula stage onwards, but restricted to either the EPI or PrE lineage upon differentiation, and late marker genes were only expressed in the mature EPI or PrE after lineage establishment.

### Lineage-specific developmental trajectories

Gene expression trajectories for each lineage were constructed by plotting the average of the normalized gene expression counts (TPM) for genes identified as lineage-specific by Petropoulos et al. ^16^, for all cells of that lineage, at each stage across the D4-7 window. The mean expression of lineage-specific genes at each day forms the trajectory. Using logarithmic regression, each GM and PM sample was fitted to the trajectory and plotted to determine an estimated “age” for each lineage for each GM and PM sample.

### Diffusion maps

Diffusion maps were generated for each lineage separately across the window of development encompassing formation of the blastocyst. Gene expression counts for genes differentially expressed in each lineage relative to all other samples (Log2FC >1, Q-value < 0.05) from the Petropoulos et al. dataset (D4-7)^16^, in combination with our GM and PM samples, were used to form diffusion maps using the scater package^81^.

### Cumulative Distribution analysis

The empirical cumulative distribution was calculated and plotted via stat_ecdf in ggplot2^79^. In each analysis, the ratio of the expression of each gene (log normalized TPM) in GM versus PM samples was calculated and plotted as a cumulative distribution. For weighted gene expression networks, this included each gene in the network (i.e., both the “from” and “to” node).

### Deconvolution of cell types within the embryonic compartment

CIBERSORT is a computational framework that can be used to quantify the proportion of cell types within a complex gene expression mixture^80^. Cell types were defined using D6-7 EPI, PrE, and polar TE samples from Petropoulos et al.^16^, and CIBERSORT was run using 1000 permutations with quantile normalization disabled. Samples undergoing deconvolution were required to have at least half of the samples with normalized read counts greater than 50, enriching for more robustly expressed genes. The relative level of each cell type within GM and PM embryonic samples, as well as in whole blastocyst samples ^32^ is presented.

### Identification of cell-cell communication networks

CellPhoneDB^41^ was used to assess cellular cross-talk between cell types of the embryo within our data and between the EPI, PrE, pTE, and mTE in the Petropoulos et al.^16^ dataset. This is done by deriving enriched ligand-receptor interactions between cell types from a repository of curated interactions, which includes both cell surface and secreted molecules. CellPhoneDB works by first permuting the cell type labels 1000 times to determine the mean expression of each ligand and each receptor within every cell type. For each ligand-receptor pair in each comparison between cell types, a null distribution is generated, representing the average expression of each ligand-receptor pair. By then calculating the proportion of the means that deviate from the mean, a p-value is calculated for the likelihood of each ligand-receptor interaction occurring in each cell type comparison. Only significant interactions (P < 0.05) were considered for all comparisons in this analysis. Resulting interaction plots were created using the chordDiagram function of the circlize (v 0.4.5) package^88^.

### Immunofluorescence analysis

Immunofluorescence was performed on an additional cohort of GM (n=3) and PM (n=3) blastocysts. As with RNA sequencing, blastocysts were frozen by vitrification on day 5/6, thawed according to standard protocols, and allowed to re-expanded in culture in G-PGD media (Vitrolife) for approximately 12 hours. The zona pellucida was removed by a very quick incubation in Acid Tyrode’s solution, followed by three washes in PBS with 0.1% BSA. Embryos were fixed with freshly prepared 4% PFA in PBS for 20 minutes at room temperature followed by three washes in PBS with 0.1% BSA and 0.1% Tween (PBST). The embryos were then permeabilized by incubation with 0.5% Triton X-100 in PBST for 20 minutes at room temperature. After permeabilization, the embryos were again washed three times in PBST and blocked with 4% normal donkey serum in PBST for 4 hours at room temperature. The embryos were incubated with primary antibodies diluted in PBST overnight at 4^0^C. Anti-CDX2 antibody (Abcam, ab76541) was used at a dilution of 1:200, anti-NANOG antibody (Millipore, MABD24) was used at a dilution of 1:200, anti- GATA4 (eBioscience (Invitrogen), 14-9980-80) was used at a dilution of 1:500, and anti-GATA-6 (R&D Systems, AF1700) was used at a dilution of 1:500). After removing the primary antibody, the embryos were washed three times in PBST and then incubated with Alexa Fluor 555 highly cross-adsorbed conjugated donkey anti-rabbit IgG (H+L) secondary antibody (Thermo Fisher Scientific, Cat. No. A-31572), Alexa Fluor 488 conjugated donkey anti-rat IgG (H+L) Highly Cross-Adsorbed secondary antibody (Thermo Fisher Scientific, Cat. No. A-21208), and Alexa

Fluor 647 conjugated donkey anti-mouse IgG (H+L) Highly Cross-Adsorbed secondary antibody (Thermo Fisher Scientific, Cat. No. PIA32787), all at a dilution of 1:500, for 1 hour at room temperature. The embryos were then washed three times with PBST and incubated with DAPI (Novus Biologicals, NBP2-31156) at a dilution of 1:500 for 30 minutes. Embryos were washed with PBST three more times before being placed in individual 2ul drops of PBS in a Nunc glass bottom dish (Thermo Fisher Scientific, 150682) and overlaid with mineral oil for imaging.

### Image acquisition

Embryos were imaged using a Nikon A1R confocal with a four-line LUN-V laser engine and DU4 detector, mounted on a Nikon Ti2 using a S Fluor 40x 0.9 NA objective. Images were acquired in resonant mode with bidirectional scanning and 4x line averaging, and 0.575 µm steps were used to collect Z-stacks of the entire embryo. The lasers used were 405nm (7% laser power), 488nm (5% laser power), 561nm (3% laser power), and 640nm (3% laser power). To avoid cross- talk between channels, Z-stacks were acquired of the DAPI and Alexa Fluor 568 channels first, and the AlexaFluor488 and Alexa Fluor 647 channels were acquired subsequently.

## Acknowledgements

We thank the generous patients who donated excess embryos to make this research possible. We also thank Miles Wilkinson, Jens Lykke-Andersen and Alon Goren for critical review of the manuscript, Joseph Owen for help with illustrations, and all members of the laboratory for helpful feedback during the project. We would like to thank the Nikon Imaging Center for help with image acquisition, analysis, and processing. This work was supported by grants from the Burroughs Wellcome Fund Career Award for Medical Scientists and the American Society for Reproductive Medicine Research Institute.

## Funding

American Society for Reproductive Medicine Research Institute (HCA, LCL, MMP)

Burroughs Wellcome Fund Career Award for Medical Scientists grant 1015559 (HCA)

Author contributions:

Conceptualization: HCA, JNC Data curation: JNC

Formal analysis: HCA, JNC, KL, KC, CT Funding Acquisition: HCA

Investigation: JNC, SS Methodology: HCA, JNC, LCL Project administration: HCA, JNC Resources: WZ, ALY, GG Software: JNC, KL, KC, CT Supervision: HCA, LCL, MMP Validation: JNC, CT Visualization: JNC, KL

Writing, original draft: HCA, JNC

Writing, review and editing: HCA, JNC, LCL, MMP

## Competing interests

The authors declare no competing interests.

## Data and materials availability

The raw sequencing data, expression-count data, and normalized read counts are deposited on NCBI GEO, with accession code GSE136106.

## Supplementary Materials

Data S1. (separate file)

Metadata for all blastocysts sequenced or stained in this study.

Data S2. (separate file)

Differential expression analysis of GM embryonic vs. GM abembryonic samples, including overlap of lineage-specific genes with Petropoulos et al., 2016^16^, and Blakeley et al., 2015^15^.

Data S3. (separate file)

Gene Set Enrichment Analysis (GSEA) for (1) GM versus PM abembryonic samples, and (2) GM versus PM embryonic samples.

Data S4. (separate file)

Identification of all ligand-receptor cell-cell communication patterns for (1) GM and PM human blastocysts and (2) single human blastocyst cells from Petropoulos et al., 2016^16^.

---

**Fig. S1.**
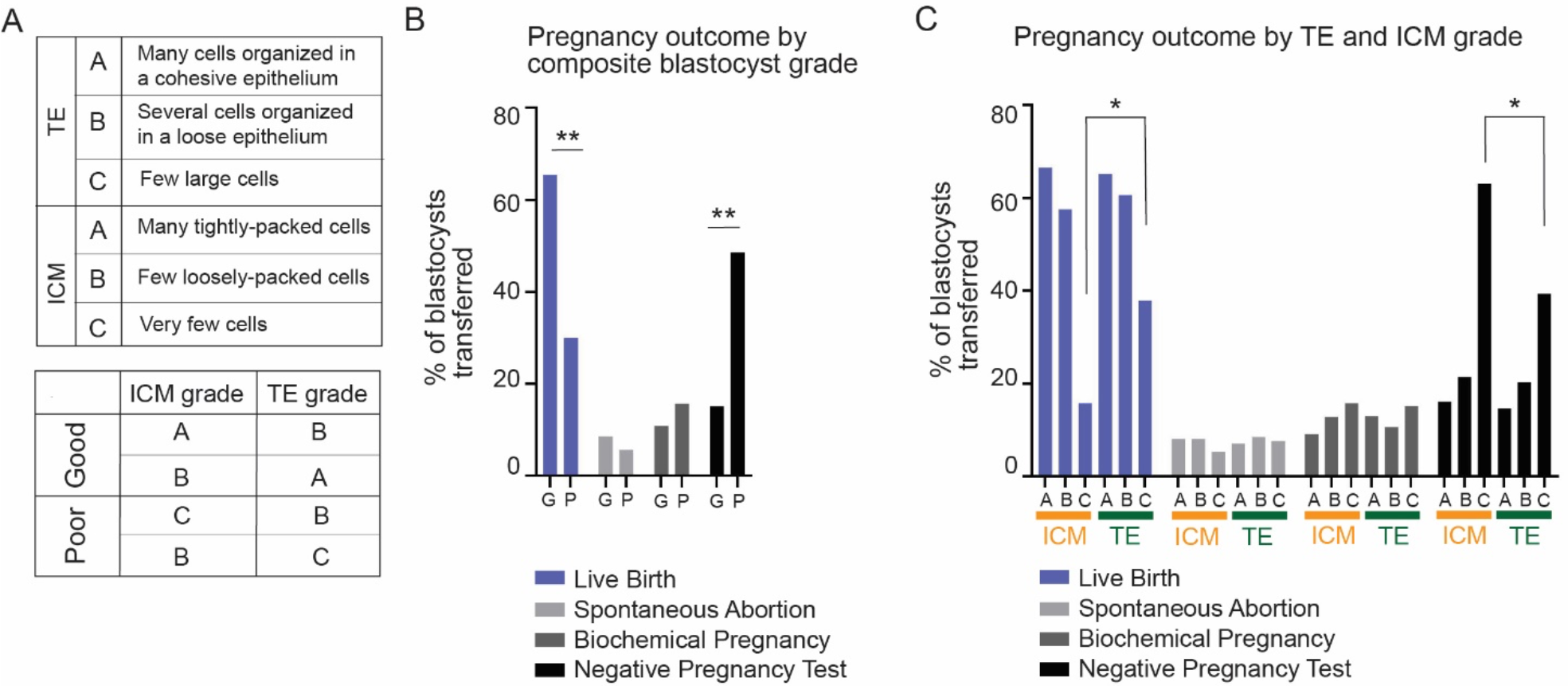
Clinical implantation is associated with blastocyst morphological grade. (A) Standard criteria used by clinical embryologists for blastocyst morphological grading used in this study^20^. **(B-C)** Retrospective clinical pregnancy outcome data following 888 euploid, single-embryo transfers in IVF cycles from 2012-2019, binned by **(B)** overall embryo grade, or by **(C)** individual ICM or TE grade (n=135, C-TE; n=27, C-ICM). A negative pregnancy test (serum β-hCG) was observed following 48.6% of PM embryo transfers compared to 15.1% for GM embryos (** p<0.001, χ2 test). Transfer of embryos with a C-graded TE resulted in a negative pregnancy test (serum β-hCG) for 39.4% of transfers compared to 63.2% of embryos with a C-graded ICM (p=0.039, χ2 test). Rates of biochemical pregnancy (15.2% C-TE vs. 15.8% C-ICM; p=0.683) and spontaneous abortion (7.6% C-TE vs. 5.3% C-ICM; p=0.618) were not different.

**Fig. S2.**
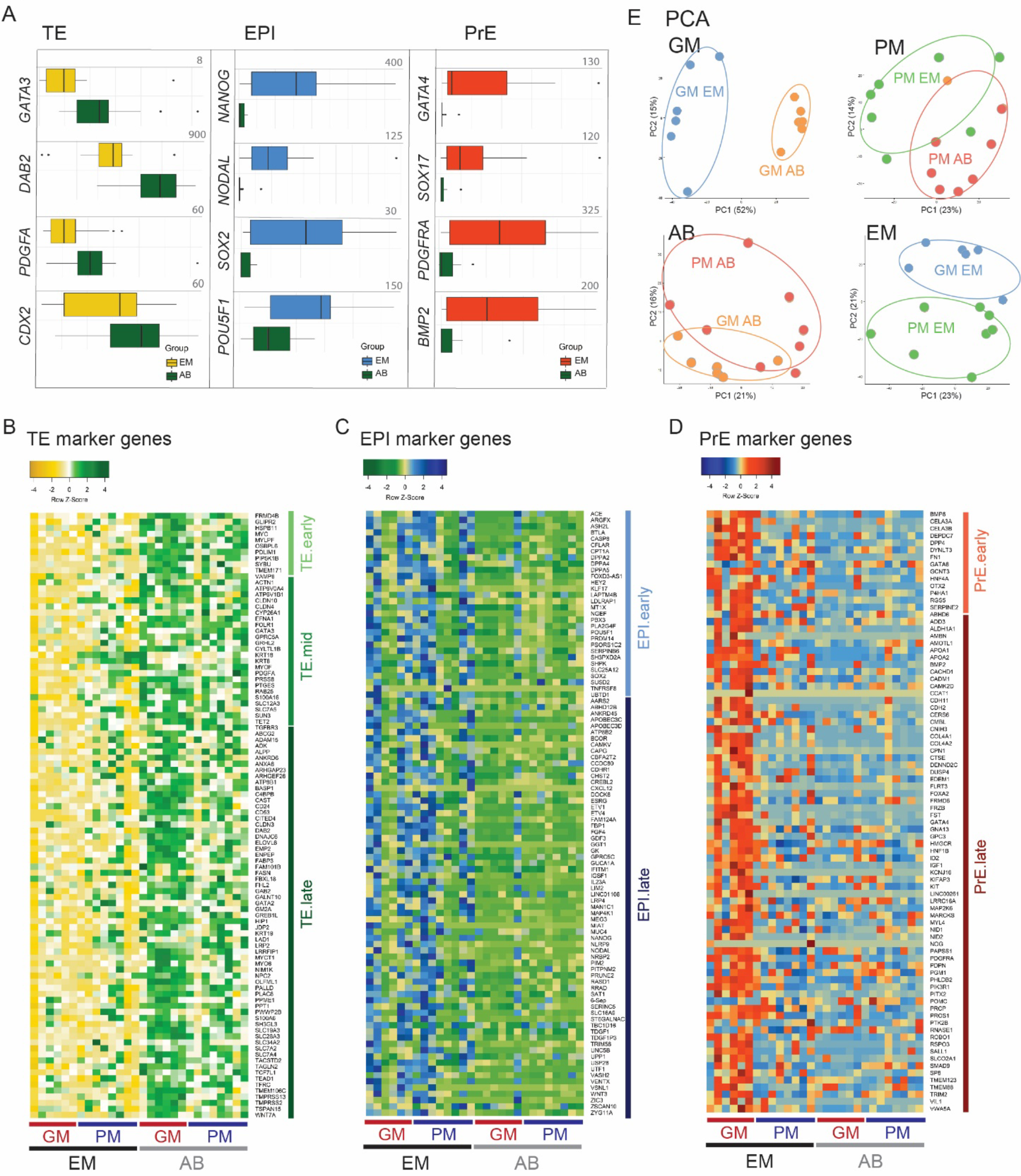
Expression of lineage markers in embryonic (EM) and abembryonic (AB) samples. (A) Box plots of normalized reads (TPM–transcripts per million) for TE, EPI, and PrE lineage markers. **(B-D)** Expression of full panels of **(B)** TE, **(C)** EPI, and **(D)** PrE lineage markers, including early, mid and late markers as previously defined^16^, with enrichment in appropriate compartments. **(E)** PCA of GM and PM samples from embryonic and abembryonic compartments.

**Fig. S3.**
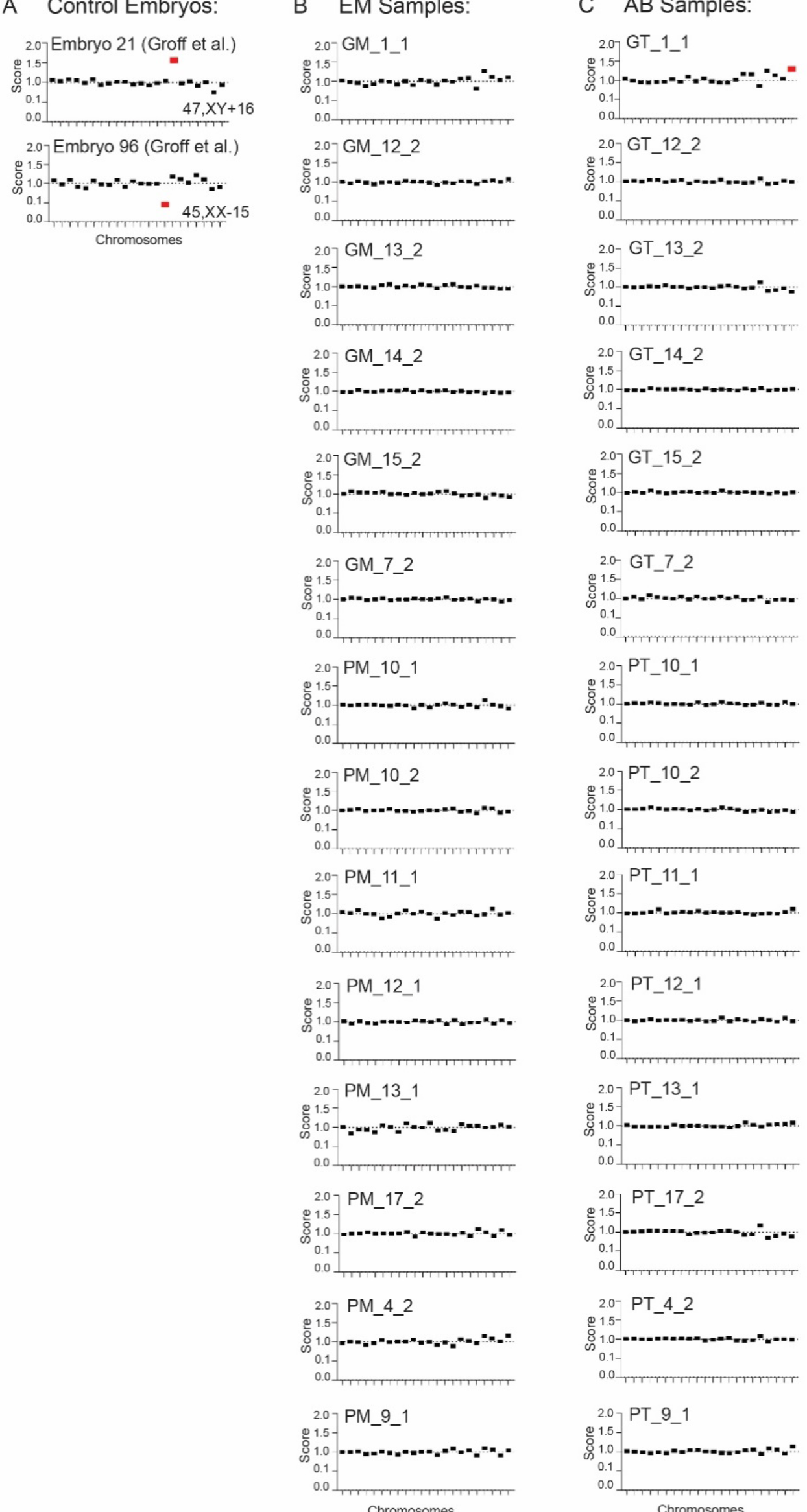
Evaluation for chromosome-wide expression imbalances in blastocyst samples. To validate PGT-A and assess the possibility of mosaicism, we examined chromosome-wide expression imbalances in our RNA sequencing data, using a published approach^24^. **(A)** First, we analyzed two published whole blastocysts^32^ as controls, confirming our ability to detect both chromosomal gains and losses. Significant aneuploid calls (Q<0.05, Chromosome Score >1.2 (Gain) or <0.8 (Loss)) are indicated in red. **(B-C)** As in (A), using this approach for our dissected **(B)** embryonic and **(C)** abembryonic samples, we were only able to detect one significant presumed mosaic aneuploidy in one GM abembryonic sample (mosaicism <30%).

**Fig. S4.**
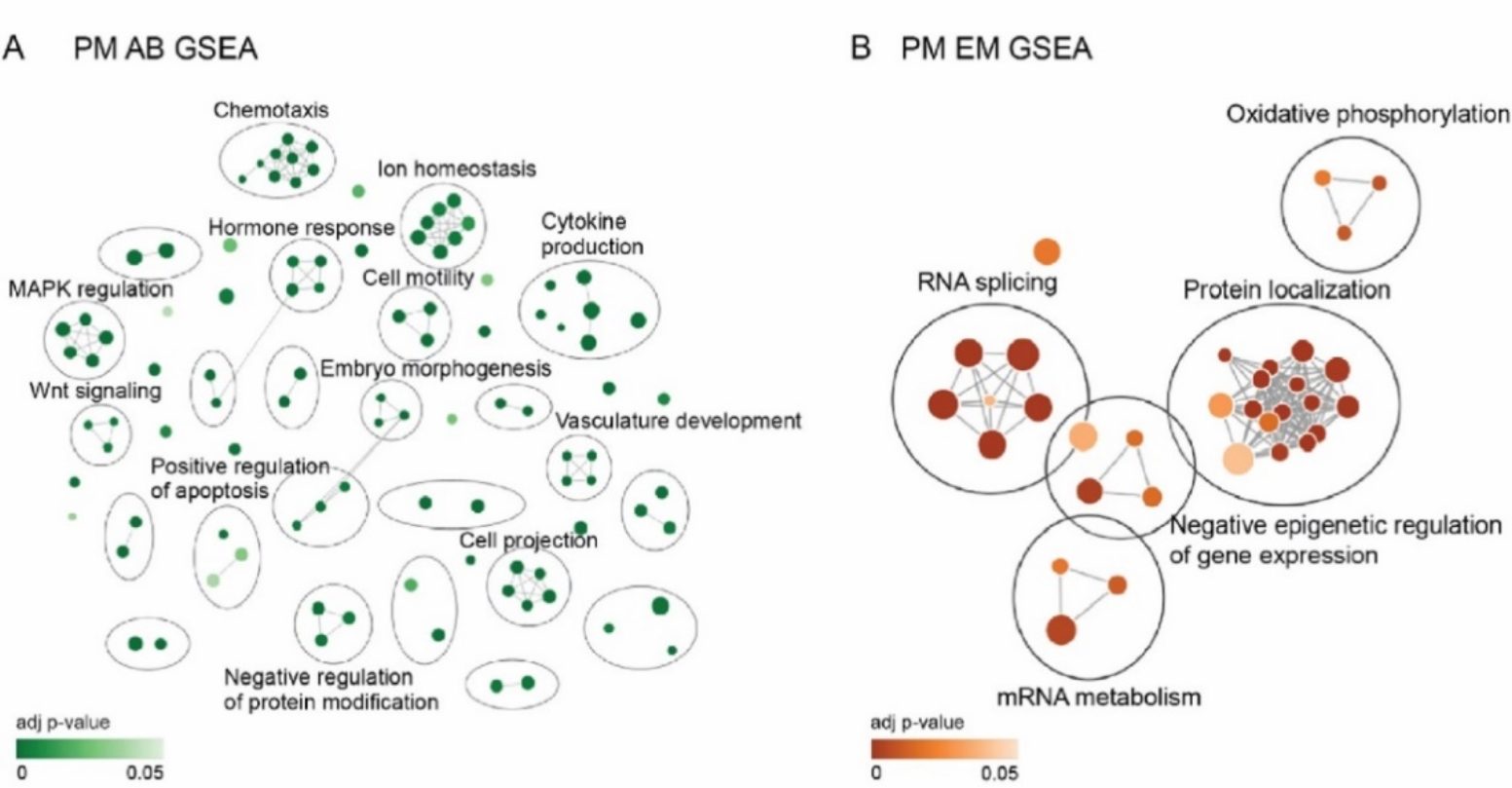
Gene ontology patterns in the abembryonic and embryonic compartments distinguish GM and PM blastocysts. Gene set enrichment analysis (GSEA) for genes most significantly upregulated (ranked by signed p-value, see Methods) in PM abembryonic (AB) **(A)** and embryonic (EM) samples **(B)**. These gene sets represent biological process gene ontologies significantly enriched (adjusted p-value < 0.05) in genes at the bottom of the ranked gene list for each embryo compartment, corresponding to genes most significantly upregulated in PM blastocysts.

**Fig. S5.**
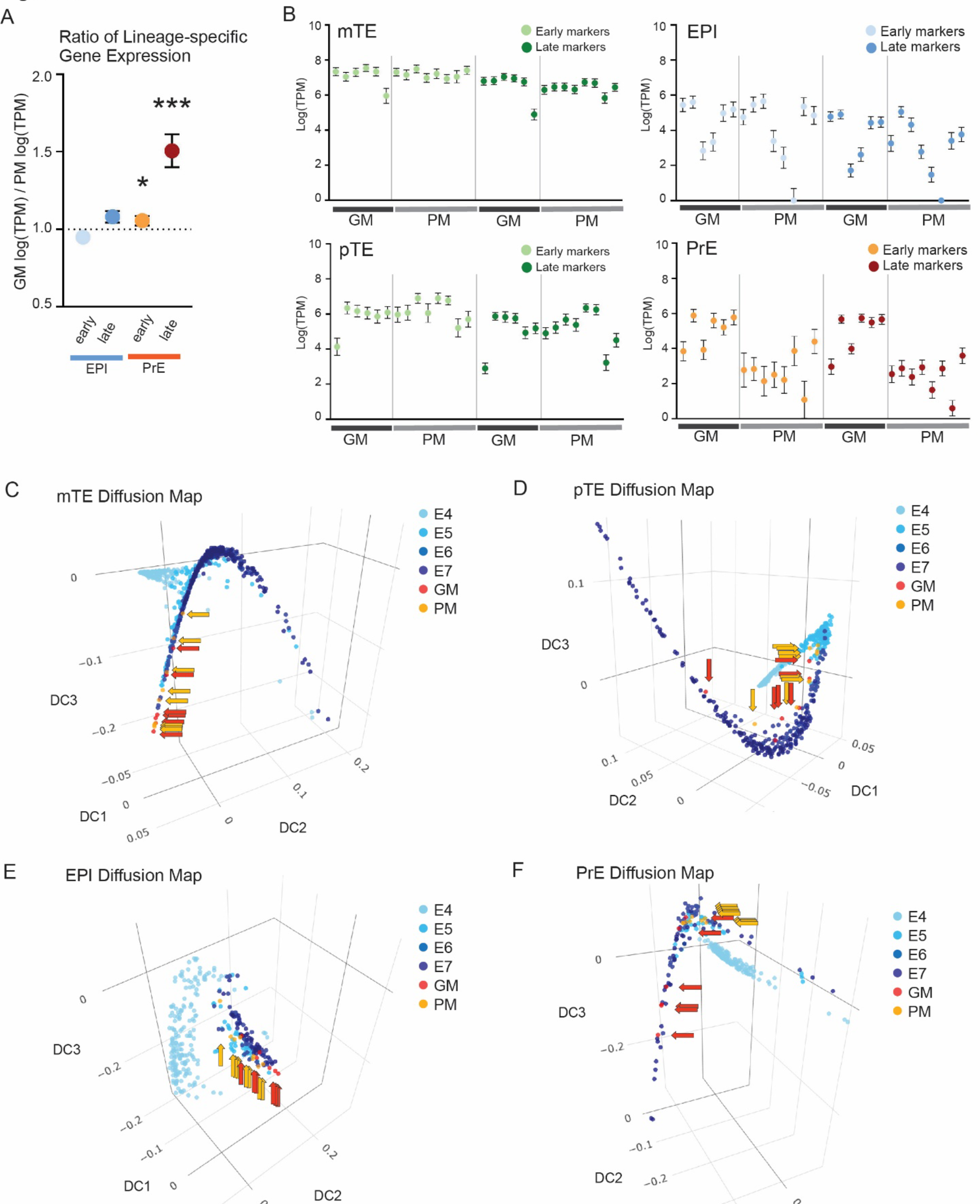
The state of PrE development is coupled with embryo morphology. (A) Ratio of the log(TPM) for early and late lineage-specific marker genes in GM versus PM blastocysts^14^. Mean and standard error indicated. **(B)** Plotted log(TPM) of early and late lineage-specific genes for individual samples, presented as mean +/- SEM. **(C-F)** Diffusion maps generated with gene expression profiles enriched in each lineage (log2FC >1, Q<0.05) demonstrating endogenous developmental progression for **(C)** mTE, **(D)** pTE, **(E)** EPI, and **(F)** PrE. For mTE, pTE, and EPI, GM and PM samples largely cluster together. However, GM samples display more advanced PrE development.

**Fig. S6.**
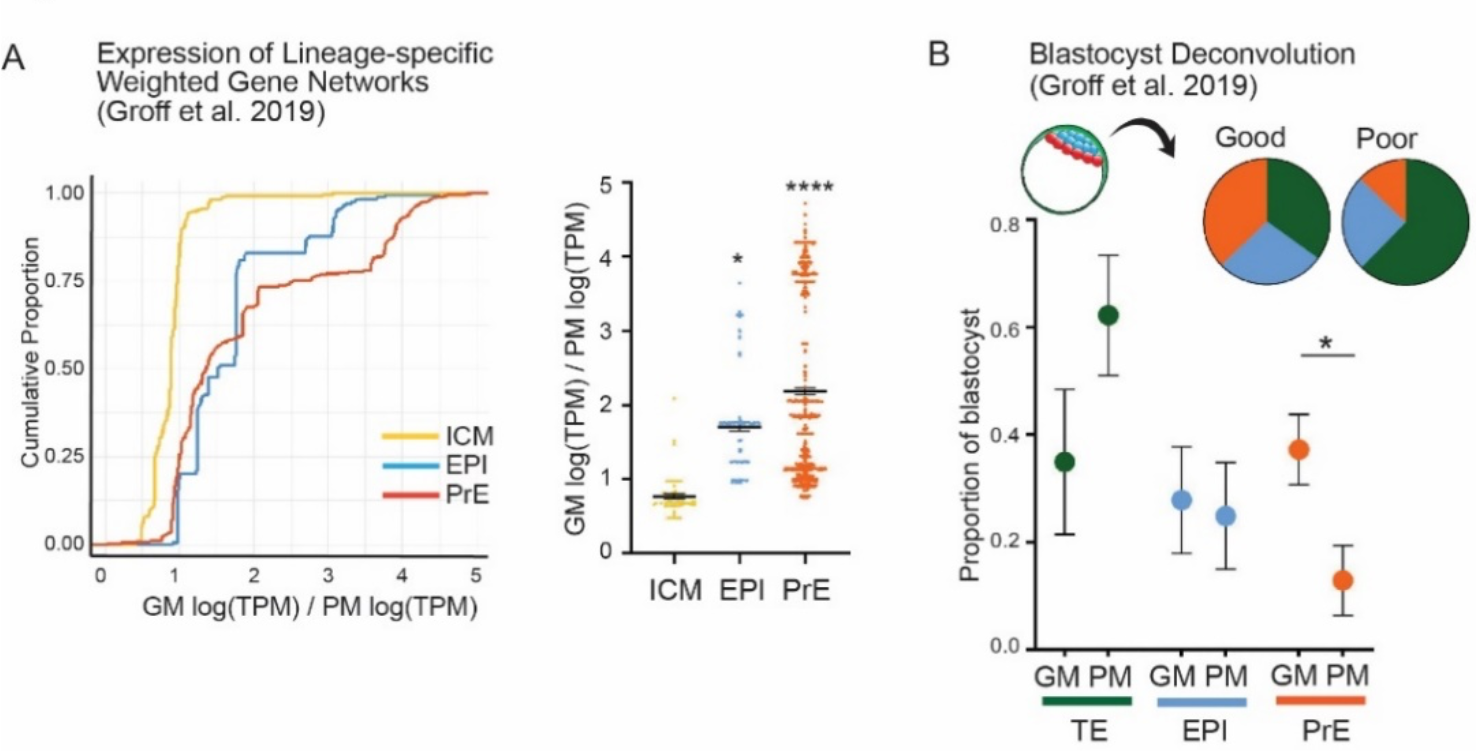
PrE defects observed in PM blastocysts are consistent across two embryo cohorts. (A) Weighted gene network cluster analysis (WGNCA) was used to define lineage-specific gene expression networks for the early, pre-differentiated ICM, EPI, and PrE lineages^14^. As in Fig 2C, the ratio of network-specific nodes in GM versus PM whole blastocysts^32^ is plotted as a cumulative distribution plot (left) and a scatter plot (right). As the expression in PM embryos decreases, the GM/PM ratio increases, and the cumulative frequency is shifted to the right, as is seen for the PrE network. **** Bonferroni-adjusted P-value = 3.9e-07 for GM versus PM PrE, * P<0.05 for GM vs PM EPI, P=1 for GM vs PM ICM, Student’s T-test. **(B)** The proportion of each lineage contributing to GM and PM whole blastocysts^32^ was predicted using a machine learning approach, as in Fig 2D^31^: GM blastocysts: 35% TE, 28% EPI, and 37% PrE; PM blastocysts: 62% TE, 25% EPI, and 13% PrE, *P<0.05, Student’s T-test.

**Fig. S7.**
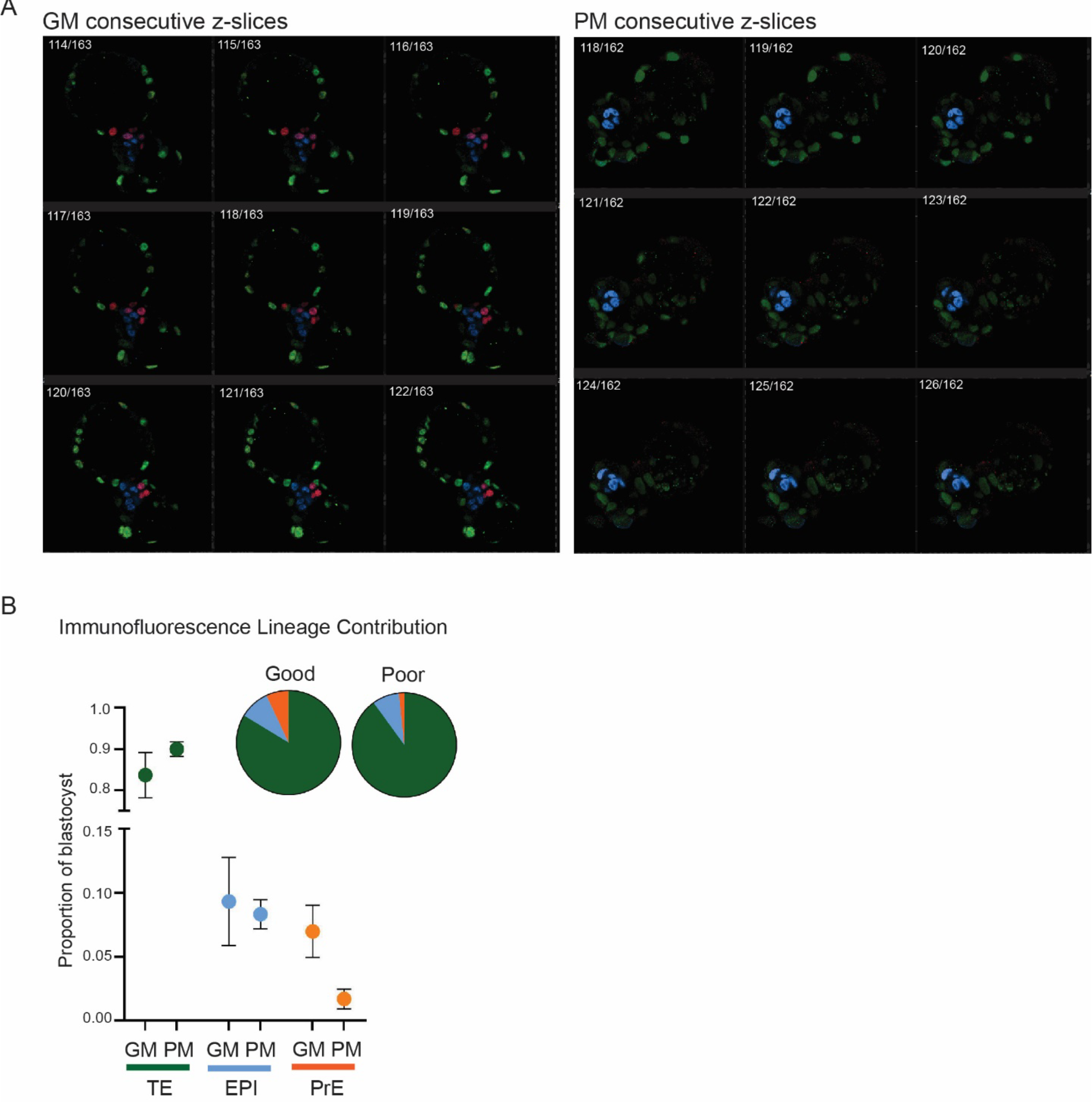
PM blastocysts contain fewer PrE cells than GM blastocysts. (A) Consecutive z-stack slices through the ICM of representative GM and PM blastocysts. Blastocysts are stained for lineage markers: CDX2 (TE), NANOG (EPI), and GATA4 (PrE). **(B)** The proportion of each lineage contributing to GM and PM whole blastocysts from immunofluorescence staining of blastocysts with markers of all three lineages, as in (A). GM (TE 85.4%, EPI 9.1%, PrE 6.9%) versus PM (TE 92.7%, EPI 8.6%, PrE 1.7%), n=3 blastocysts per group. Corresponding cell count data is demonstrated in Fig 3C.

**Fig. S8.**
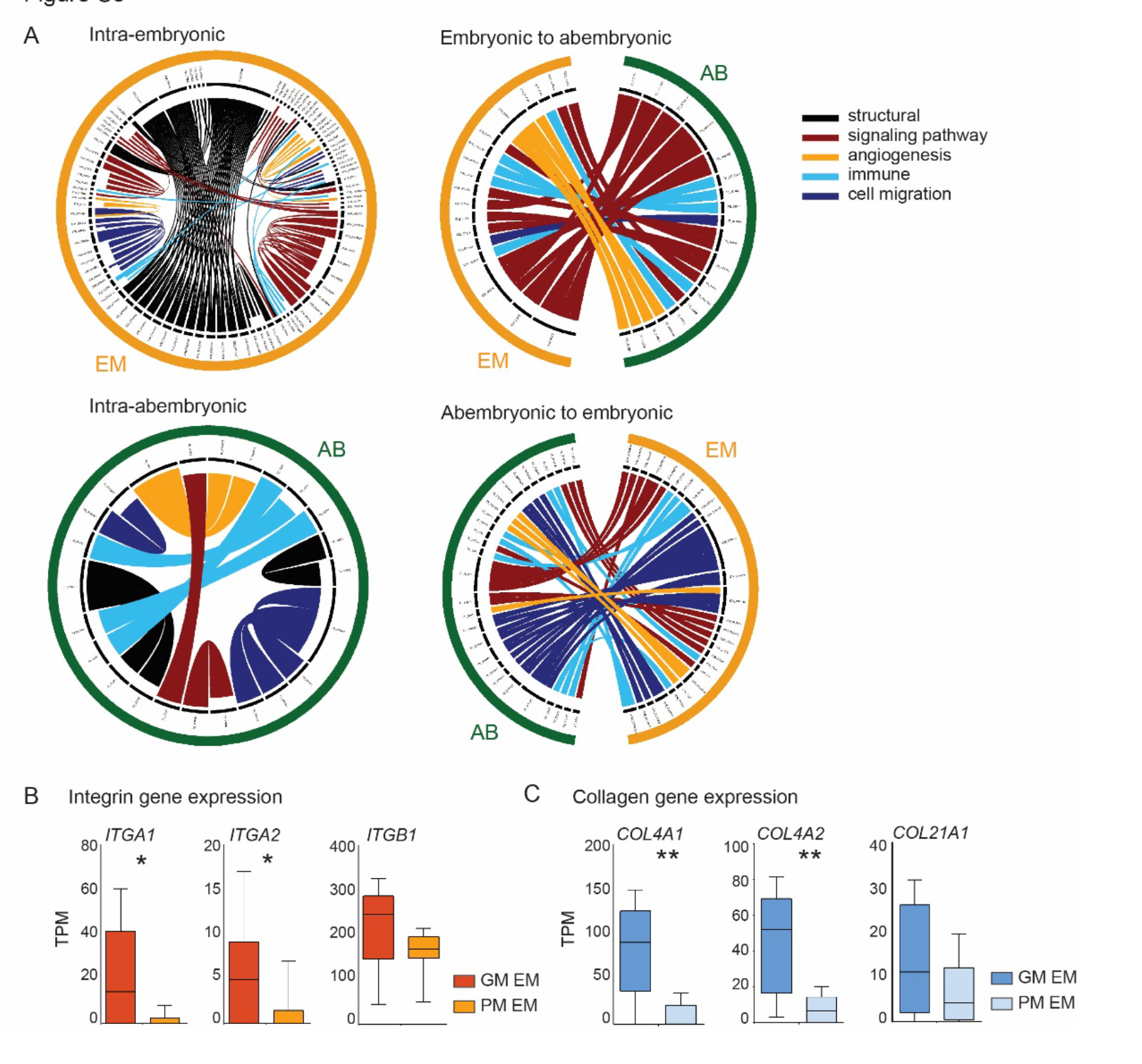
Cell-cell communication networks predicted in GM blastocysts. (A) Interaction categories predicted in GM blastocysts (Intra-embryonic, Embryonic-to-abembryonic, Abembryonic-to-embryonic, and Intra-abembryonic) include “structural”, “signaling pathway”, “angiogenesis”, “immune”, and “cell migration”, colored as defined in the key. **(B-C)** Normalized counts (TPM) for the most enriched PrE-associated **(B)** integrin and **(C)** collagen subunits in PM versus GM embryonic samples. Median and standard error indicated, **P-adj<0.005, *P-adj<0.05.

**Fig. S9.**
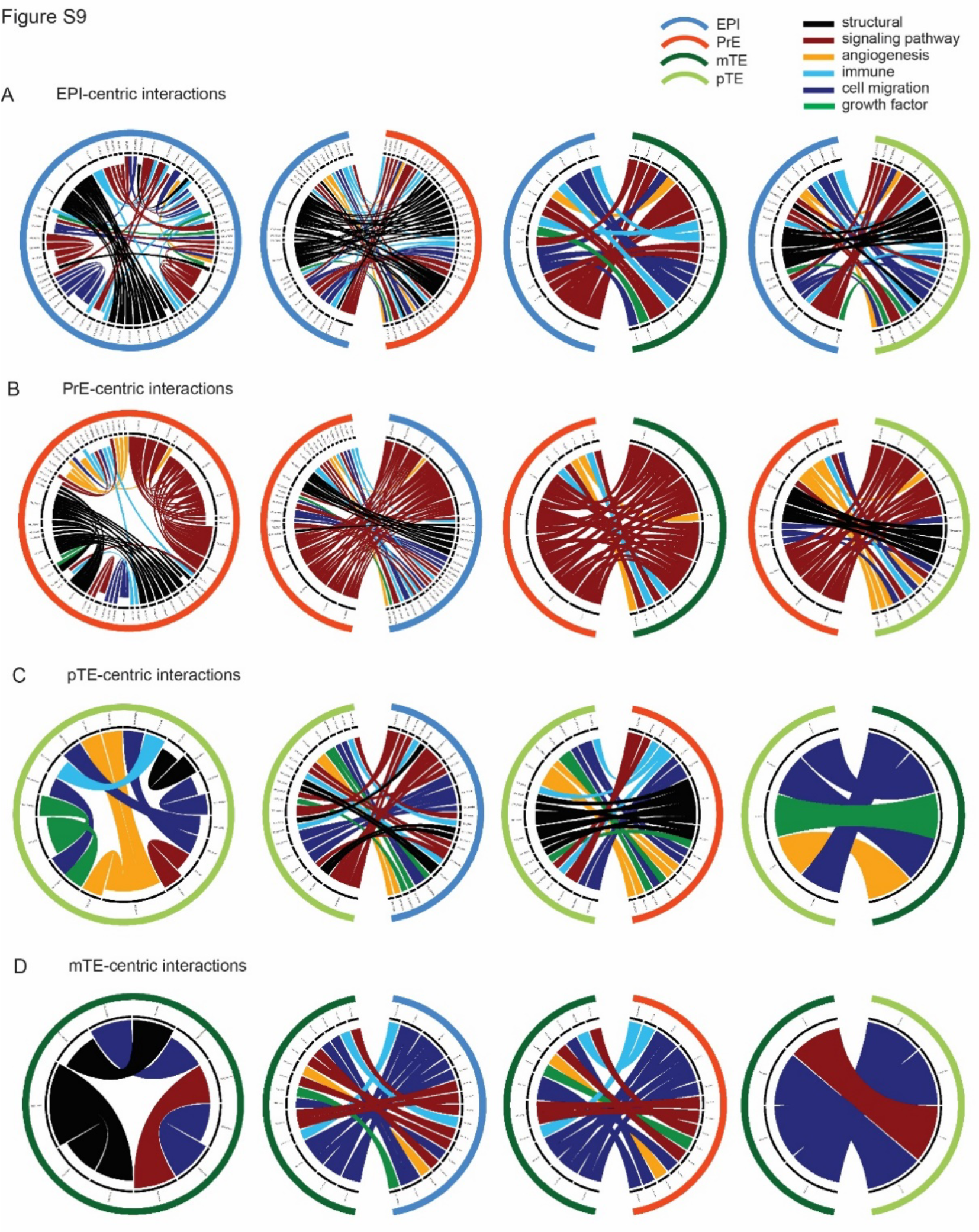
Lineage-specific cell-cell communication networks predicted in the blastocyst. (A-D) Categories of interactions identified in D5-7 human blastocysts include “structural”, “signaling pathway”, “angiogenesis”, “immune”, “cell migration”, and “growth factors”, colored as defined in the key. **(A)** EPI-centric interactions include intra-EPI signals, as well as signals originating in the EPI and communicating with PrE, pTE, and mTE, with EPI labeled as blue on the exterior rim. **(B)** PrE-centric (orange), **(C)** pTE-centric (light green), and **(D)** mTE-centric signals (dark green), as in (A). A complete resource of lineage-specific interactions predicted in the blastocyst can be found in Table S4.

**Fig. S10.**
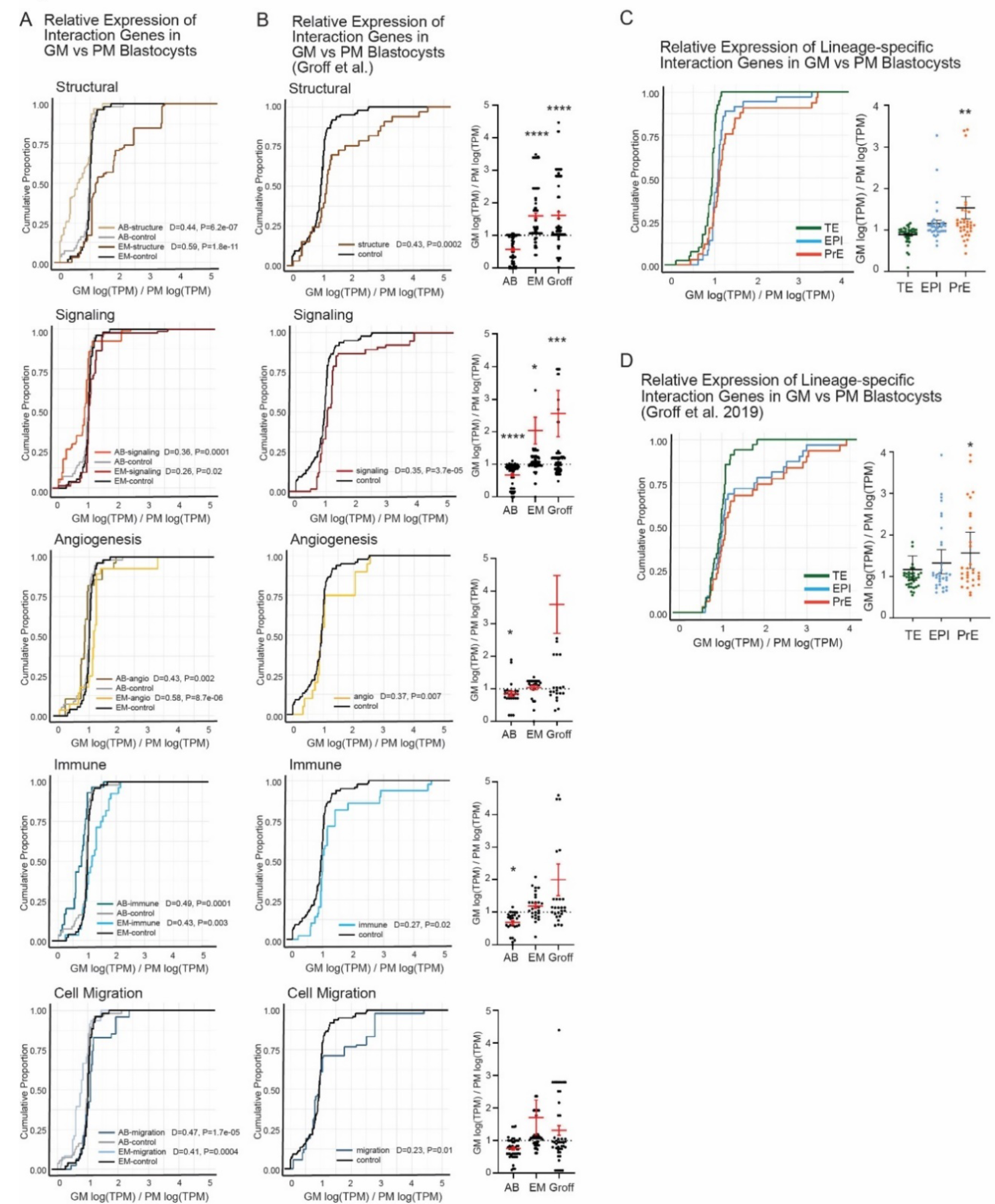
PrE-specific and structural communication networks are enriched in GM embryos and in an independent embryo cohort. (A-B) The relative expression of ligand-receptor genes in GM versus PM **(A)** embryonic (EM) and abembryonic (AB) samples and **(B)** whole blastocyst samples is presented as the cumulative distribution. A group of randomly chosen genes is also plotted as a control (n=100). The relative (ratio of) expression of ligand-receptor genes is plotted as a scatter plot for comparison. *** Bonferroni-adjusted P-value <0.001, **** Bonferroni-adjusted P-value <0.0001. **(C)** Lineage-specific communication as defined in this study (Fig S9)^16^, in which the gene product initiating communication (i.e., the “from” node) is assayed. The ratio of the expression of these lineage-specific communication genes initiated in the TE, EPI, and PrE is presented here as a cumulative distribution plot (left) and scatter plot (right) of the ratio of the expression in GM versus PM blastocysts. ** P<.01. **(E)** As in (A), but for whole blastocyst samples^32^. * P=0.06.

## Notes

### Competing Interest Statement

The authors have declared no competing interest.

